# Functional interaction between *Drosophila* olfactory sensory neurons and their support cells

**DOI:** 10.1101/2021.10.04.463011

**Authors:** Sinisa Prelic, Venkatesh Pal Mahadevan, Vignesh Venkateswaran, Sofia Lavista-Llanos, Bill S. Hansson, Dieter Wicher

**Author notes:** Corresponding author: PD Dr. Dieter Wicher.

## Abstract

Insects detect volatile chemicals using antennae, which house a vast variety of olfactory sensory neurons (OSNs) that innervate hair-like structures called sensilla where odor detection takes place. In addition to OSNs, the antenna also hosts various support cell types. These include the triad of trichogen, tormogen and thecogen support cells that lie adjacent to their respective OSNs. The arrangement of OSN supporting cells occurs stereotypically for all sensilla and is widely conserved in evolution. While insect chemosensory neurons have received considerable attention, little is known about the functional significance of the cells that support them. For instance, it remains unknown whether support cells play an active role in odor detection, or only passively contribute to homeostasis, e.g. by maintaining sensillum lymph composition. To investigate the functional interaction between OSNs and support cells, we used optical and electrophysiological approaches in *Drosophila*. First, we characterized the distribution of various supporting cells using genetic markers. By means of an *ex vivo* antennal preparation and genetically-encoded Ca^2+^ and K^+^ indicators, we then studied the activation of these auxiliary cells during odor presentation in adult flies. We observed acute responses and distinct differences in Ca^2+^ and K^+^ fluxes between support cell types. Finally, we observed alterations in OSN responses upon thecogen cell ablation in mature adults. Upon inducible ablation of thecogen cells, we notice a gain in mechanical responsiveness to mechanical stimulations during single-sensillum recording, but a lack of change to neuronal resting activity. Taken together, these results demonstrate that support cells play a more active and responsive role during odor processing than previously thought. Our observations thus reveal that support cells functionally interact with OSNs and may be important for the extraordinary ability of insect olfactory systems to dynamically and sensitively discriminate between odors in the turbulent sensory landscape of insect flight.

## 1. Introduction

Olfaction is an ancient and critical sensory modality for all animals. Sensitivity to volatile chemicals underpins a great variety of essential behaviors for survival and reproduction such as foraging for food, avoidance of biotic and abiotic hazards, sexual mating, reception of inter- and intraspecific semiochemicals (Vosshall, 2000), and is a both ubiquitous and principal sense for metazoan life (Ache and Young, 2005). The perception of airborne cues begins by the detection of odors by dedicated, specialized sensory organs. Though the astounding variety of smelling organs may seem diverse, the general features of olfactory systems are conserved and share several invariable features which allow for specific and sensitive sampling of broad ranges of odors (Eisthen, 1997; Krieger and Breer, 1999; Ache and Young, 2005; Eisthen and Polese, 2007; Ng et al., 2020). It has long been noted that even disparate olfactory tissues such as mammalian olfactory mucosa and arthropod sensilla display striking similarities in olfactory transduction and structural constraint (Shirsat and Siddiqi, 1993; Abbas and Vinberg, 2021). With respect to cellular repertoire, olfactory organs are always composed of odorant receptor-equipped sensory neurons innervating an epithelium, and a lesser-explored set of auxiliary cells that co-arise in development, which remain closely apposed to their corresponding neurons, and are thought to play roles in maintaining and potentiating the ability of neurons to perform their sensory function (Schmidt and Benton, 2020).

The various populations of support cell types and the functions of these “support networks” have been partially elucidated across myriad organisms, which indicate that these cells fulfill many hitherto unknown or understated tasks that many be endemic to sensory systems across disparate organisms (Charlton-Perkins et al., 2017). For instance, mounting evidence points to the important role of many support cells in regulating sensory neuronal activity, transmission and structural integrity. In the mouse auditory system, cochlear support cells reduce neuron cell excitability by modulating extracellular space and the speed of K^+^ redistribution through osmotic shrinkage (Babola et al., 2020). Ommatidial cone support cells in the *Drosophila* compound eye functionally interact with photoreceptor neurons through means of altered metabolism and ion homeostasis (Charlton-Perkins et al., 2017). In the *C. elegans* peripheral chemosensory system, the amphid sheath glial cell (AMsh) is able to autonomously respond to aversive chemicals and consequently suppresses its amphid ASH neuron through GABA release to promote olfactory adaption (Duan et al., 2020). In *C. elegans* mechanosensors, nose touch receptors crucial for touch behaviors depend on ion homeostasis performed by supporting glial cells harboring Na^+^/K^+^ ATPases (Johnson et al., 2020). Support cells have also garnered much attention following the COVID-19 pandemic (Cooper et al., 2020), with studies revealing the non-neuronal expression of SARS-CoV-2 entry genes in the sustentacular support cells of mammalian olfactory systems, which are implicated as central players in the symptomatic anosmia following infection (Brann et al., 2020). Even though the role and influence of support cells in different modalities and organisms is beginning to be uncovered, a large gap in our understanding remains. For instance, what differentiates adjacent support cell types? To what degree, and on what temporal scale are support cells involved in the sensory process? Do animals show physiological or behavioral differences contingent on the variability in support cell phenotype? And though sensory systems show varying degrees of conservation and anatomical parallelisms, such as neuronal compartmentalization (Ng et al., 2020), which functional elements beside neurons are selected or free to vary, and which remain stable in evolution?

Particularly across the range of insect taxa, sensory systems are generally conserved, and take the form of sensilla, chitinized hair-like protrusions from the cuticle on insect bodies, most often acting in chemo-, mechano-, hygro- and thermo-sensing with similar underlying cytological organization (Steinbrecht, 1996; Chai et al., 2019). Sensilla of the insect model organism *Drosophila melanogaster* are typically innervated by one-or-few sensory neurons, and are classified based on morphological shape as well as identity of sensory neurons that innervate them. These sensory neurons individually express distinct receptors from a wide range of receptor families such as the odorant receptor (OR), gustatory receptor (GR), ionotropic receptor (IR), pickpocket (Ppk) and transient receptor potential (TRP) protein families (Gallio et al., 2011; Joseph and Carlson, 2015), as well as non-canonical transporter-receptors recently described such as Amt (Vulpe et al., 2021). *Drosophila* specifically possess many appendages with chemosensory sensilla, such as maxillary palps, proboscis, wings, sexual organs (e.g. ovipositor), bodily bristles, and tarsi. However, the most particularly tractable and broadly studied system is that of the antenna, a bilaterally-occurring appendage on insect heads. *D. melanogaster* antennae are segmented into three parts, named scape, pedicel and funiculus, referring to first, second and third antennal segments, respectively. The funiculus is characterized by an arista, a large modified bristle that arises from the proximal part of the funiculus and extends laterally. The segment also houses the largest collection of olfactory sensilla in *Drosophila*.

This olfactory system in particular presents an amenable model to study cellular and molecular underpinnings of olfaction. To date, concerted efforts have led to a near-complete description of the antennal architecture, namely, an anatomical atlas of the distribution and identification of all sensillum types, as well as an exhaustive understanding of the number, identity and odor-tuning properties of ORs and their respective olfactory sensory neurons (OSNs) innervating each sensillum (Montell, 2021). Standardized electrophysiology techniques such as single-sensillum recording, along with expansive, genetic toolkits for genetic manipulation by virtue of binary expression systems, such as the GAL4/UAS, LexA/LexAop and Q binary expression systems (Potter et al., 2010; Riabinina and Potter, 2016) and contingent neurogenetic methods, such as the delta-halo empty neuron system (Hallem et al., 2004), have permitted numerous investigations into understanding the interface between the external odor world and the receptors and OSNs, which process and ultimately transmits odor information to the brain (Grabe and Sachse, 2017). To date, functional studies utilizing *Drosophila* antennae have revealed important, far-reaching molecular and ecological insights and exposed conserved principles in peripheral coding and odor information processing. However, the majority of *Drosophila* antennal cells, namely support cells, have received considerably less attention and stand in stark juxtaposition to neurons with respect to functional characterization. It remains unclear whether this family of sensory support cells play active or passive roles in sensing both internal and external cues, whether they are coupled or synchronized with local neuronal activity, and whether they mobilize responses to maintain and potentiate *Drosophila* sensory neuron activity to achieve robust odor detection.

For instance, what influence or role do these perineuronal support cells have on odor responsiveness and odor processing? In recent years, few hints have surfaced. They are thought to play a passive role in lymph maintenance by releasing various ‘helper’ proteins into the sensillum lymph to aid in stimulus recognition (Rihani et al., 2021). Among these are the odor-degrading enzymes (ODEs), a poorly-defined set of catabolic enzymes which play a role in odor degradation and clearance (Younus et al., 2014; Baldwin et al., 2021), and chemosensory proteins (CSPs) which act as extracellular holdase chaperones in their ability to bind and transport chemicals in aqueous environments (Pelosi et al., 2018; Zhu et al., 2019). In insect olfactory tissues, the best studied accessory proteins are the odor-binding proteins (OBPs), which have long been posited to influence odor responsiveness by allowing less soluble odors to traverse the lymph and interact with olfactory receptors. Indeed, OBPs have often been shown *in vitro* to complex with odorants and affect OR selectivity and sensitivity (Wang et al., 2020). The combinatorial expression of OBPs by support cell type and by sensillum type has been shown through fluorescent *in situ* RNA hybridization (Larter et al., 2016). Curiously, follow-up studies have demonstrated that the removal of OBPs, including the most abundant transcripts in antennal transcriptomes, seems to have no impact on odor responses in basiconic sensilla, with the singular exception of Obp19a Obp28a double mutant responses to one compound, linoleic acid (Xiao et al., 2019). It is a striking observation that OBPs seem to some degree dispensable for olfaction *in vivo*, given that RNA-seq screens reveal OBPs are the most highly expressed genes in the antenna (Menuz et al., 2014). Recent evidence in *Helicoverpa armigera* suggests that pheromone-specific OBPs can contribute to peripheral sensitivity to pheromones, but play negligible roles in modulating response kinetics as well as selectivity of ORs *in vivo* (Guo et al., 2021). Separately, support cells have also been shown to be essential for the correct biogenesis of functional sensilla (Andrés et al., 2014; Ando et al., 2019). Support cells have likewise been shown to express sensory neuron membrane proteins (SNMPs) alongside neurons in various insect species and may play important roles in odor detection as membrane-spanning molecules involved in the capture, transport, clearance or presentation of odorants to ORs (Li et al., 2014; Cassau and Krieger, 2020). More recently, the non-canonical olfactory transceptor and ammonium transporter Amt, has been shown to be widely expressed in *Drosophila* antennal support cells (Vulpe et al., 2021), a first indication that these non-neuronal cells are equipped with nutrient or odorant sensing elements, a phenomenon known to occur widely beyond sensory neurons (Holsbeeks et al., 2004). Despite this, the apparent diversity of auxiliary cells, and the extent of their participation in odor reception and transduction remains largely unaddressed. This is surprising, given widespread conservation of sensillum cell architecture across insect genera (Steinbrecht, 1996), the shared terminal differentiation and developmental origin with sensory neurons (Ghysen and Dambly-Chaudiere, 1989; Hartenstein and Posakony, 1989, 1990; Chai et al., 2019), as well as the structural homology with various other non-chemosensory organs such as chordotonal organ scolopidia and the arista (Keil and Steinbrecht, 1984; Foelix et al., 1989; Yack, 2004). Moreover, auxiliary cells form numerous seals and contacts around the sensory neuron by virtue of septate junctions, as well as maintain closed compartments such as the sensillum lymph and perineuronal lumen between neuron and thecogen cell (Steinbrecht, 1980; Keil and Steinbrecht, 1983, 1987; De Kramer, 1985; Shanbhag et al., 2000). Lastly, auxiliary cells display clear morphological specializations, such as the rich presence of microvilli, proteasomes, mitochondria and vesicles (Shanbhag et al., 2000; Nava Gonzales et al., 2021), suggesting that auxiliary cells function and retain continuous metabolic activity throughout insect adulthood.

The repertoire of *Drosophila* support cells occurs in a highly stereotyped fashion. Aside from sensory neuron(s), whose axons may be enveloped by one or more glial cells, each sensillum is populated by a thecogen, tormogen and trichogen cell. These have been termed sheath, socket and shaft/bristle/hair cells, respectively, and are collectively referred to as accessory, auxiliary or supporting cells. Though largely treated as functionally equivalent due to their poor molecular characterization and an inability to experimentally manipulate them separately from one another, the triad of support cell types has been dissociated partially based on physiological identity and unique features evident during development. For instance, tormogen cells in particular have been reported to express specific cytochrome P450 enzymes at their apical poles (Willingham and Keil, 2004) and display features of active transport across the apical portion, which is covered in microvilli. In *Calliphora* blow flies, this apical portion features an enrichment in mitochondria, particles and vesicles, and exhibits non-specific alkaline phosphatase and Mg^2+^-activated ATPase activity (Gnatzy and Weber, 1978). In the silk moth, two tormogen cells arise during development, where the inner tormogen cell degenerates within 2 days post-apolysis (Keil and Steiner, 1990, 1991). Atypically to the general rule of one support cell type per sensillum, *Drosophila* coeloconic sensilla are populated with two tormogen cells (Shanbhag et al., 2000). A mature tormogen cell is distinct from the trichogen cell in that it extends a characteristically long, stalk-like protrusion that terminates below the level of the OSN soma, at least in *Drosophila* (Nava Gonzales et al., 2021). Though the validity of the marking is somewhat unclear, tormogen cells have been historically tagged using a promoter found upstream of the Suppressor of Hairless gene (Su(H)) termed ASE5 (Barolo et al., 2000), a DNA-binding protein component of the Notch signaling pathway (Bray and Furriols, 2001) which reportedly contributes to the terminal differentiation of the tormogen socket and trichogen shaft cells (Schweisguth and Posakony, 1992, 1994; Gho et al., 1996). Its counterpart, the trichogen cell, is uniquely involved in the olfaction-essential formation of nanopores on the sensillum surface by modifying the cuticular envelope during metamorphosis in an Osiris gene-dependent fashion (Ando et al., 2019). Moreover, in *Drosophila* embryo chemosensory organs, the trichogen cell has been shown to expresses artichoke (atk), a gene required for correct morphogenesis of ciliated sensory organs like sensilla, without which larvae exhibit impaired chemotaxis (Andrés et al., 2014). Currently, no adult cell type-specific markers are known to mark the trichogen cell. As a result, studies have been limited in their ability to dissociate tormogen cells from trichogen cells without the use of the high resolution approach of electron microscopy. Finally, thecogen cells are characterized by their tight, innermost association with their OSN(s). Their plasma membranes are closely apposed, whereby the thecogen cell envelops the soma and inner dendrites of the neuron, terminating basally below the soma and apically at the root of the sensillum shaft (Keil, 1997). In practice, thecogen cells are therefore often loosely termed glial cells, due to their sheath-like morphology, and incorrectly confused with olfactory ensheathing (glial) cells which arise - as well as function more prominently - at the antennal lobe, which migrate into the antennal periphery to insulate OSN axon fibers (Sen et al., 2005; Wu et al., 2017). In cockroach maxillary palps, thecogen cells have been observed to bear lysosomes, perform endocytosis, and have been suggested to clear the lymph space of odorant stimuli (Seidl, 1992). Thecogen cells have been shown to express nompA, a structural protein implicated in tethering neuronal dendrites to the sensillum shaft (Chung et al., 2001). nompA mutants are therefore anosmic due to the broad inability of OSNs to attach and innervate into sensilla (Chung et al., 2001).

Interestingly, all three support cell types occur broadly across both extrasensory and chordotonal sensilla (Chung et al., 2001; Yack, 2004; Kamikouchi et al., 2009; Göpfert and Hennig, 2016). For example, each scolopidium in the Johnston’s organ is comprised of a triad of non-neuronal cells: a ligament cell, a scolopale cell, and a cap cell, which are equivalent to the triad found in olfactory tissues. Though named differently depending on which organ they are found in, this collection of supporting cells is both ubiquitous across sensory modalities, disparate body parts and insect genera, which suggests these cells perform vital roles with regard to sensory perception (Kaissling, 1986; Schmidt and Benton, 2020).

In this study, we characterize all known ways to access these cells for experimental manipulation by systematically exploring the distribution and reliability of specific cell markers of the heterogeneous types of non-neuronal cells in the antenna. Subsequently, we use live cation imaging in an *ex vivo* antennal preparation to characterize whether major supporting cell classes detectably respond to odor presentation events, and find concomitant ion fluxes immediately following neuronal activation by the odor proxy and synthetic agonist VUAA1. We observe ion-specific physiological differences between thecogen and tormogen auxiliary cell responses, which indicate that these cells are functionally distinct yet both coupled to OSN activity. Last, by way of *in vivo* single sensillum recording (SSR) between adult flies with intact and ablated thecogen cells, we find differences in responses to a panel of ecologically relevant odorants, without change to OSN resting activity in flies lacking thecogen cells. Curiously, we find a gain in mechanosensitivity in thecogen cell-free flies, as well as hints of sensillum-specific effects. Given a long research history but relative scarcity of insight, we also speculate on the potential action of support cell types with respect to the structure and function of insect olfactory tissues. Altogether, our live cation imaging and electrophysiological examinations indicate that support cells acutely respond to chemical cues or neuronal activity in real time, and can no longer be viewed as passive or stimulus-acquiescent elements of the *Drosophila* olfactory system. This renewed consideration may have broad implications for our understanding of odor processing in insect sensory apparatuses within and beyond chemosensory sensilla, as well as complex multicellular sensory compartments in other species.

## 2 Materials & Methods

### Lines used

A list of transgenic fly lines used in this study can be found in Table 1. This study on the vinegar fly *Drosophila melanogaster* was conducted in Germany where research on invertebrates does not require a permit from a committee that approves animal research. The transgenic fly laboratory meets all requirements of the Thuringian State Office for Consumer Protection (verbraucherschutz.thueringen.de).

**Table 1.** List of transgenic fly lines used.

| Genotype | Source | ID | Where used (*purpose*) |
| --- | --- | --- | --- |
| UAS-mCD8::GFP | Okada et al., 2009 |  | immunohistochemistry (*membrane-localizing GFP*) |
| UAS-mCD8::RFP | BDSC | 27399 | immunohistochemistry (*membrane-localizing GFP*) |
| nompA::GFP | BDSC | 42694 | immunolabeling (*of nompA protein*) |
| UAS-rpr | BDSC | 5824 | inducible apoptotic knockout of thecogen cells (*apoptosis-inducing protein*) |
| αTub84B-GAL80<sup>ts</sup> | BDSC | 7019 | inducible apoptotic knockout of thecogen cells (*drives pancellular temperature-dependent GAL4 repression*) |
| UAS-GCaMP6f | BDSC | 42747 | cation imaging (*genetically encoded Ca<sup>2+</sup> indicator*) |
| Orco-GAL4 | BDSC | 26818 | cation imaging (*drives expression in Orco<sup>+</sup> OSNs*) |
| nompA-GAL4 | C.M. (UCSB) |  | immunohistochemistry, cation imaging, inducible apoptotic knockout of thecogen cells (*drives expression in thecogen cells*) |
| ASE5-GAL4 | C.M. (UCSB) |  | immunohistochemistry, cation imaging (*drives expression in tormogen cells*) |
| repo-GAL4 | BDSC | 7415 | immunohistochemistry (*drives expression in Drosophila glia*) |
| elav-GAL4 | BDSC | 8760 | immunohistochemistry (*drives pan-neuronal expression*) |
| Orco-lexA | Root et al., 2008 |  | immunohistochemistry (*co-staining in parallel with GAL4/UAS, drives expression in Orco<sup>+</sup> OSNs*) |
| UAS-mCD8::RFP; lexAop-mCD8::GFP | BDSC | 32229 | immunohistochemistry (*parallel fluorescence labeling when driven by GAL4 and lexA constructs*) |
| UAS-GINKO1 | Custom-made |  | cation imaging (*custom line; genetically encoded K<sup>+</sup> indicator*) |
| UAS-GAL4 | Hassan et al., 2000 |  | immunohistochemistry (*used to constitutively drive expression*) |
| yw; CyO/Bl; TM2/TM6B | BDSC | 3704 | balancing and crossing |

### Fly stocks

All fly lines were obtained from the Bloomington Drosophila Stock Center, Indiana (bdsc.indiana.edu) except where otherwise noted. ASE5-GAL4 and nompA-GAL4 lines were kindly provided by Craig Montell (University of California, Santa Barbara). The ASE5-GAL4 enhancer fragment was originally constructed from an *in vivo* expression assay using *lacZ* reporter gene analysis; ASE5 refers to a 372bp 5′ subfragment of an enhancer containing five S*uppressor of Hairless (Su(H))* binding sites, termed “ASE5”, that drove high and specific expression in tormogen (socket) cells in a variety of adult bristles (Barolo et al., 2000). The exact extent of the enhancer fragment used to construct nompA-GAL4 is to our knowledge unknown but has originally been used to drive expression in scolopale cells of the Johnston’s organ and is derived from the promoter of no mechanoreceptor potential A (*nompA)* gene (Chung et al., 2001; Todi et al., 2004, 2005).

### DNA vector construction of pUASTattB-GINKO1 vector and generation of UAS-GINKO1 flies

The genetically-encoded fluorescent potassium indicator GINKO1 (Shen et al., 2019) was prepared from DNA sequence kindly provided by Yi Shen, and inserted into plasmids to prepare pUAST-GINKO1. The resulting constructs were sequenced, amplified by PCR and purified conventionally. Subsequent *D. melanogaster* germline transformation with the prepared plasmid was carried out by BestGene using the PhiC31 integration + Cre-loxP removal plan (Bestgene, www.thebestgene.com). The vector was inserted into chromosome III to produce the genotype *+;+;UAS-GINKO1/TM3*. Flies were acclimatized to local rearing conditions for several generations prior to crossing.

### Fly rearing

All *D. melanogaster* flies were maintained on conventional cornmeal agar medium (recipe available in data repository) in incubation under a 12h/12h light/dark cycle at 25°C and 70% humidity. For heatshock experiments inducing apoptotic cell death of thecogen cells in adult flies, flies were crossed and subsequently reared entirely at 18°C from lain egg onward, and heatshocked at the GAL80^ts^-permissive temperature of 32°C for 24h or 48h, to induce apoptosis by way of lifting GAL80 repression of the GAL4 transcription factor.

### Open antenna preparation

Antennae of 2-12 day old flies were excised and prepared as described previously (Mukunda et al., 2014; Halty-deLeon et al., 2018). Briefly, flies were anesthetized on ice. Antennae were excised using a fine needle, deposited into a 100 μl droplet of *Drosophila* Ringer solution (5 mM HEPES; 130 mM NaCl; 5 mM KCl; 2 mM MgCl_2_; 2 mM CaCl_2_; 36 mM sucrose), equilibrated to pH = 7.30 and room temperature. Excised antennae were then fixed in vertical position with a two-component silicone curing gel (KWIK-SIL, World Precision Instruments, https://www.wpi-europe.com). Thereafter the funiculus was cut horizontally with micro-scissors, exposing a layer of antennal tissue, and immersed in an additional 800 μl Ringer solution for immediate imaging. All antennae were immersed in solution for the duration of the imaging experiments. All experiments were carried out during the day (light cycle).

Where variable KCl concentrations were used experimentally, Ringer solutions were prepared afresh by aliquoting KCl-free Ringer solution into separate bottles, and anhydrous KCl (Carl Roth, carlroth.com) was added to reach the following final KCl concentrations per bottle: 1, 3, 10, 100, and 150 mM KCl. Prior to use, all Ringer solutions were equilibrated to room temperature and pH = 7.30.

### Immunohistochemistry

For whole antennal mount preparations, female flies between 4 days and 8 days old were collected. Antennae were dissected into a solution of Ringer with 0.1% Triton X-100 (PT solution). The tube was spun down using a benchtop centrifuge and solution was siphoned and replaced with 4% paraformaldehyde (PFA) in PT solution for 2h on ice on a shaker. Subsequently, the antennae were washed 4x by replacing existing medium with fresh PT solution for 20 minutes between each wash. Antennae were then blocked for 1h at 20°C with 10% bovine serum albumin in PT solution, or 5% normal goat serum (NGS) in PT solution where goat antibodies were to be used (blocking solution). For immunostaining, primary antibodies were then incubated with the sample in blocking solution for 48h at 4°C. Antennae were then washed as before and blocked once more with blocking solution for 2h at 20°C prior to secondary antibody incubation. For secondary antibody incubations, samples were incubated overnight at 4°C in the dark on a shaker. Antennae were finally washed 4x with PT solution and mounted onto slides in Vectashield (Vector Laboratories) for confocal imaging.

Antennal sections were prepared by depositing dissected antennae into OCT Mounting medium for cryotomy (VWR, vwr.com) and frozen at -80°C. Cryotomy was performed by plating 10 μm cryosections of dissected antennae deposited onto adherent glass slides using a Microm HM 560 Cryostat (Thermo Scientific). Sections were immediately fixed in 4% PFA in phosphate-buffered saline (PBS) for 10 minutes, and washed gently 2x for 10 minutes in PBS. For permeabilization and blocking, sections were permeabilized in PBS with 0.1% Triton X-100 (PT buffer) for 30 minutes, and then transferred to a humidified chamber for blocking with blocking solution (PTS buffer: PT buffer + 5% NGS) for 30 minutes. 100 μl of primary antibody solution in PTS buffer was pipetted onto the slide and incubated overnight at 4°C. Next, the slides were washed 3x for 10 minutes using PT buffer on a shaker and blocked in PTS buffer for 30 minutes. Slides were subsequently incubated with secondary antibodies for 2h at room temperature in the dark. Slides were washed 3x for 5 minutes using PT buffer, and finally mounted using 60 μl Vectashield (Vector, Burlingame, CA, USA) under a coverslip.

For staining nompA>RFP & nompA::GFP or Orco>GFP, we used the primary antibodies chicken anti-GFP and rabbit anti-RFP (Invitrogen, Carlsbad, CA, USA), at the relative dilutions of 1:1000 and 1:500, respectively, and the secondary antibodies goat anti-chicken-A488 and goat anti-rabbit-A546 (Invitrogen) at the relative dilutions of 1:250 and 1:500, respectively. For staining nompA>UAS-GAL4>GFP, we used the primary antibodies rabbit anti-RFP 1:1000 (Invitrogen), and the secondary antibodies goat anti-rabbit 1:100 or 1:250 (Invitrogen). For all DNA stainings, we used 1:1000 Hoechst dye staining.

### Confocal microscopy

Micrographs were captured using a cLSM 880 (Carl Zeiss, Oberkochen, Germany) using 10×, 20×, 40× or 63× water immersion objectives (C-Apochromat, NA: 1.2, Carl Zeiss). Where Airyscan is noted, images were obtained using the Airyscan detector and mode on the cLSM 880 (Huff, 2015). Where linear unmixing is noted, images were obtained using linear unmixing mode in ZEN software (Carl Zeiss, www.zeiss.com). Z-stack maximum intensity projection images were obtained at 1 μm intervals for whole antennal overviews and 0.5 μm intervals for detailed sections and close-ups. All confocal images were adjusted for contrast and brightness with ZEN software (Carl Zeiss).

### Live cation imaging

Ca^2+^ and K^+^ imaging was performed with an epifluorescence microscope (Axioskop FS, Zeiss, Jena, Germany) coupled to a monochromator (Polychrome V, Till Photonics, Munich, Germany). A water immersion objective (LUMPFL 40× W/IR/0.8; Olympus, Hamburg, Germany) was used along with an imaging control unit (ICU, Till Photonics). A 490 nm dichroic mirror and a 515 nm long-pass filter were employed to filter emitted light for capture with a cooled CCD camera controlled by TILLVision 4.5.62 software (TILL Photonics). An experimental protocol was programmed to sample images every 5 seconds over 180 cycles, allowing for 15 minutes of continuous specimen imaging. Each sampling event follows a 50 ms exposure to 475 nm light generated by the monochromator. All chemical applications to the sample were performed by pipetting a volume of 100 μl of 100 μM VUAA1 and/or 100 μM CdCl_2_ in Ringer solution onto the immersed objective for advection and diffusion over the submerged antenna. Afterward, a background region was marked along with observed cells, which were marked as regions of interest (ROIs). Where ROIs were subsetted for parallel analysis, each region was qualitatively judged as responding (responder) or non-responding (non-responder) to the treatment based on whether a change in signal was noticeable upon visual inspection, and were labeled as such for subsequent processing. TILLVision software was used to generate a matrix of average fluorescence values for background region and ROIs; this matrix was exported for data analysis using *R*.

### Data analysis and visualization of cation imaging experiments

Cation imaging response magnitudes were calculated as average changes in ROI fluorescence signal subtracted from background signal, relative to a non-response window of time of 10 imaging cycles and converted into percentage change relative to baseline (ΔF/F_0_), as used previously (Mukunda et al., 2014; Halty-deLeon et al., 2018). A custom script was written in *R* to transform the exported matrix of raw fluorescence intensity values into ΔF/F_0_ time course plots for each of the regions of interests as marked on the open antenna. The script reads a batch of replicates to produce a final time course plot showing all replicates and an average with its standard error of the mean (SEM) for each time point. First, background noise (an ROI with an area outside the antenna in the image) is subtracted from all ROIs in each antenna (replicate) for background noise-correction. Second, each time course is normalized to a baseline of 0 based on a common ‘resting’ time window (10 imaging cycles over 50 seconds) prior to the first stimulation, so that biological replicates can be compared. Lastly, a mean average and standard error of the mean is calculated for each time point across all replicates (individual antennae) or across type of ROI (responder vs. non-responder ROI). The script produces two outputs here: a table of processed data (for purposes of statistical analysis), and a time course graph. Here, the calculated average time course is plotted superimposed on its source replicates to show both individual and grouped average trends, along with labels demarcating time points at which treatments occurred during imaging, and a gray-shaded interval showing the time window used for normalizing each ROI recording (F_0_ time-window). For concentration curves, the time point post-stimulation with VUAA1 used was selected by determining the time at which a maximum or minimum response is reached across all replicates and treatments. All error bars represent SEMs. Paired and unpaired Student’s *t*-tests were used to compare two sets of data. Two-way ANOVA with the Tukey *post hoc* tests were used to compare multiple sets of data. Asterisks indicate statistical significance (^n.s.^p>0.05, *p≤0.05, **p≤0.01, ***p≤0.001, ****p≤0.0001). Statistical analyses were performed using GraphPad Prism 4-9 (www.graphpad.com), Rstudio 1.3-1.4 (www.rstudio.com) and Microsoft Excel.

### Electrophysiology (SSR)

Single sensillum recordings were performed on *D. melanogaster* antennae. A 2+ day old fly was held immobile in a 200 μl pipette tip and fixed on a glass side with laboratory wax. The third antennal segment was fixed in such position that the medial-posterior side faced the observer. Extracellular recordings were done using electrochemically (3 M KOH) sharpened tungsten electrodes by inserting ground electrode into the eye and recording electrode into the base of sensilla using a micromanipulator system (Luigs & Neumann SM-10). Sensilla were visualized with 1000× magnification using a binocular microscope (Olympus BX51WI). Signals were amplified (Syntech Uni-versal AC/DC Probe, www.syntech.nl), sampled (96000/s) and filtered (3kHz high-300Hz low, 50/60 Hz suppression) using a USB-IDAC. Neuronal activity was recorded using AutoSpike software (v3.7) for 3 seconds pre- and 10 seconds post-stimulus. Stimuli were delivered for 500 ms and were added to pre-humidified air being constantly delivered onto the fly at a rate of 0.6 LPM. Stimuli were prepared by pipetting 10 μl of desired compound, dissolved in hexane (10^−4^), onto a filter paper of diameter 10 mm. No more than 5 sensilla were recorded from each fly and odors were used a maximum of 5 times with a gap of 30 minutes for re-equilibrating pipette headspace.

### Quantitative analysis of peak frequency responses and area-under-curve of treatment responses

Response frequency plots were generated in AutoSpike by selecting recordings and creating frequency time courses using 25 ms bins. All time courses were saved with a timestamp of when odorant presentations were applied, such that they could be aligned in post-processing. Response frequency plots were charted using GraphPad Prism 9.0.1 based on *n*=4-9 flies for each of the 16 treatments, 3 fly genotypes, and 2 heatshock conditions. For air gust corrections, we used a three-step pipeline. We initially plot the raw response trace (step 1). The mean response trace for air gust treatments was then subtracted from other traces, and the resulting time courses and their mean traces are shown labeled as ‘air gust corrected’ (step 2). These average-corrected traces were then smoothed with a 400 ms rolling average (step 3) using the *rollmean* function within the *zoo R* package to (i) remove leftover response artefacts due to micro-timing mismatches in stimulus onsets left over from subtracting the air gust responses previously, and (ii) to leave only large effects behind that would more easily be judged qualitatively. A 400 ms duration for rolling average was selected beforehand based on typical response durations of approximately 400 ms, which we deemed a conservative (response-removing) approach. All resulting traces of the three-step processing workflow were subsequently mined for peak (maximum) response and area-under-curve using a custom *R* script specified to only survey data points within a predefined ‘response window’ of stimulus onset to 1 second after stimulus onset, with the exception of the smoothed data where the ‘response window’ was defined as 0.5 seconds prior to stimulus onset to 1 second after stimulus onset, due to the shifting of the responses by the smoothing transformation. Area-under-curve calculations were achieved using the *trapz* trapezoidal integration function in the *pracma* package in *R*. For sensillum response profiles, we performed two-way ANOVA with the Tukey *post hoc* tests to compare multiple sets of data across genotype and heat treatment. Asterisks indicate statistical significance, as before.

### Chemicals

All chemicals used for SSR electrophysiology were purchased from Sigma (www.sigmaaldrich.com) with the highest purity. Chemicals used were selected for being diagnostic odors for each of the individual neurons of the ab1, ab2, ab3, ab4 and ab5 sensillum subtypes and for the ab6A neuron (total of 13 chemicals). The compound, the neuron that it is a diagnostic odor (best ligand) of, and the CAS number or source follow: ethyl acetate, ab1A, 141-78-6; ethyl lactate, ab2A, 97-64-3; CO_2_, ab1C, human exhalation; methyl salicylate, ab1D, 119-36-8; methyl acetate, ab2A, 79-20-9; ethyl-3-hydroxybutyrate, ab2B, 5405-41-4; ethyl hexanoate, ab3A, 123-66-0; 2-heptanone, ab3B, 110-43-0; E2-hexanal, ab4A, 6728-26-3; geosmin, ab4B, 16423-19-1; geranyl acetate, ab5A, 105-87-3; pentyl acetate, ab5B, 628-63-7; 1-octen-3-ol, ab6A, 3391-86-4. VUAA1 (N-(4-ethylphenyl)-2-((4-ethyl-5-(3-pyridinyl)-4H-1,2,4-triazol-3-yl)thio)acetamide) was synthesized by the Mass Spectrometry/Proteomics group at the Max Planck Institute for Chemical Ecology (Jena, Germany). VUAA1 was dissolved in DMSO to yield a 100 mM stock solution. CdCl_2_ solution was prepared by adding anhydrous CdCl_2_ (Sigma) to yield a stock of 100 mM CdCl_2_ in Ringer solution as described prior. In cation imaging experiments, VUAA1 and CdCl_2_ stock solutions were solved 1:1000 in the same Ringer solution on the day of the experiment to yield fresh 100 μM solutions for application onto the antennal specimen during live cation imaging.

## 3. Results

### Evaluation of *Drosophila* support cell markers for the third antennal segment

The third antennal segment of *D. melanogaster* is populated by a plethora of neuronal and non-neuronal cell types, including a discrete set of auxiliary cells, which form stereotypical multicellular arrangements and compartments sealing the sensillum lymph and enveloping the OSNs which project their dendrites, often branched, into the lymph space for chemical sensing (Figure 1A). More distantly from the OSN are epithelial cells that flank sensillum compartments entirely, and glial cells that insulate OSN axons projecting into the olfactory lobe via an olfactory nerve (Figure 1A). Sensilla cover the entire surface of the funiculus, including the protruding arista and sacculus pits, which also host the same cellular repertoire as sensilla (Figure 1B). Given the scarcity of characterization of non-neuronal funiculus cell markers, we began by exploring and validating all available ways to label each support cell type by use of *Drosophila* binary expression systems (Viktorinová and Wimmer, 2007). Broadly, we looked to systematically examine expression patterns and suitability as marking devices.

**Figure 1.**
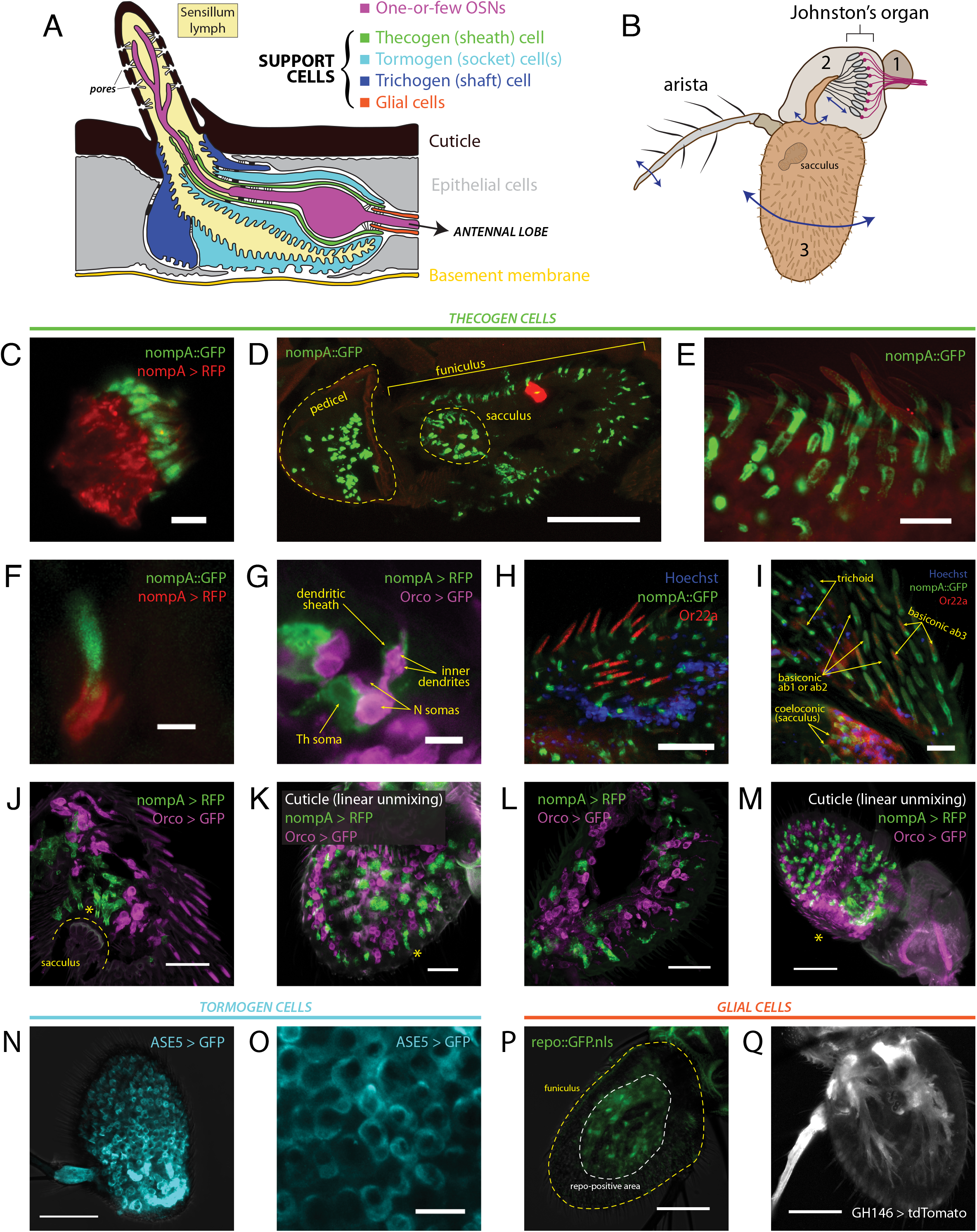
Heterogeneity of antennal cells: patterns of expression of various support cell markers and cell type-specific drivers. (**A**) Diagram of a typical insect olfactory sensillum and its constituent structures and cells. (**B**) Illustration of an entire *Drosophila* antenna. Antennal segments are numbered. Vibrational/rotational movements of arista and segment 3 (funiculus) are transduced by the Johnston’s organ in segment 2 (pedicel), while segment 3 and its characteristic pit (sacculus) harbor the vast majority of chemosensory sensilla on the antenna. (**C**) Immunofluorescence of scolopale cells and nompA protein in the Johnston’s organ. Scale bar: 5 μm. (**D**) Immunofluorescence micrograph of longitudinally cryosectioned *D. melanogaster* antenna, showing presence of nompA protein in both scolopale cells of the pedicel and cells in the funiculus and sacculus. Scale bar: 50 μm. (**E**) Maximum intensity projection of confocal imaging of whole antenna using Airyscan. nompA protein forms distinct tube-like, sheathing structures under sensilla. Scale bar: 10 μm. (**F**) Immunofluorescence of sectioned antenna showing thecogen cells excreting nompA extracellularly, similarly to scolopale cells in the second antennal segment. Scale bar: 2 μm. (**G**) nompA GAL4-driven reporter expression marks thecogen cells, visibly ensheathing sensory neurons past the inner dendrite in a sectioned antenna. Scale bar: 5 μm. (**H**) Max intensity projection of Airyscan imaging of whole antenna showing thecogen cells marked with nompA protein co-localizing in sensilla positive for OR22a, indicating thecogen presence in ab3 sensillum subtype. Nuclei are stained with Hoechst fluorescent dye. Scale bar: 20 μm. (**I**) Thecogen cells marked with nompA are present in all sensilla types. Immunofluorescence indicates thecogen cell presence in a variety of basiconics (in both OR22a-positive ab3 and OR22a-negative ab1 or ab2 sensilla), in trichoid sensilla, and in coeloconic sensilla of the sacculus. Max intensity projection of Airyscan imaging of whole antenna. Scale bar: 10 μm. (**J**) Thecogen cell expression driven by nompA-GAL4 shows presence of thecogen cells in sacculus, in Orco-negative sensilla. Max intensity projection of Airyscan imaging of whole antenna. Asterisk denotes position of sacculus sensilla. Scale bar: 20 μm. (**K**) Whole mount of antenna following linear unmixing for cuticular auto-fluorescence. Asterisk denotes a sensillum containing a thecogen cell without an attributable Orco-labeled OSN. Scale bar: 20 μm. (**L**) Antennal section showing OSNs without attributable sheathing thecogen cells. Scale bar: 20 μm. (**M**) Whole mount linear unmixing for cuticular auto-fluorescence. Asterisk denotes the proximomedial area, opposite the arista, without any nompA-GAL4 labeled thecogen cells. Scale bar: 50 μm. (**N**) Max intensity projection of whole antenna showing ASE5-GAL4 driving ubiquitous and uniform expression of a GFP reporter in the third antennal segment. It labels more globular-shaped cells that are more distal from the cuticle than thecogen cells. ASE5-GAL4 is purported to mark tormogen cells. Scale bar: 50 μm. (**O**) Digital magnification of micrograph shown in panel O. Marked cells are well tessellated and of similar morphology upon inspection. Scale bar: 10 μm. (**P**) The *Drosophila* glial cell marker repo does not mark any cells superficial to the cuticle; white dashed line indicates extent of repo signal, yellow dashed line outlines the antennal edge. Scale bar: 50 μm. (**Q**) Max intensity projection of whole antenna showing glial marking driven by GH146-GAL4. Glial cell bodies are distant from sensilla, similar to repo stainings. Scale bar: 50 μm.

First, we performed immunofluorescence co-imaging of both nompA protein and nompA promoter-driven expression of a RFP fluorescent reporter, to check whether protein and promoter reliably labeled the same cells. Due to a notable homology between chordotonal organs and olfactory sensilla, we were aware that scolopale cells and thecogen cells share the expression of nompA (Chung et al., 2001). Thus, we looked at the second antennal segment (pedicel), where nompA-GAL4 and nompA protein have been previously used to label scolopale cells and the mechanical linkage of the scolopidia to the cuticle making up the Johnston’s organ (Chung et al., 2001; Roy et al., 2013). As expected, we found nompA-GAL4 drove expression in scolopale cells, which were positive for GFP-tagged nompA (Figure 1C). Next, we looked at the localization of nompA protein across a longitudinal cross-section of both second and third antennal segments, finding nompA protein across the entirety of the funiculus, including within the characteristic sacculus pits (Figure 1D). Upon closer inspection with super-resolution imaging, clear tube-like sheathing structures of nompA protein in all observed sensilla were apparent (Figure 1E). To better understand whether the protein localization conformed or overlapped with thecogen cellular shape, we found that the nompA-GAL4 driven cytoplasmic reporter expression did not overlap with that of nompA protein (Figure 1F), suggesting that nompA is excreted or exported from cells with an active nompA promoter, and that nompA protein is not suitable as a cellular marker. We additionally checked whether nompA is found within the apical sensillum lumen, for the possibility that it would colocalize with Orco or coat the outer dendrites. We found nompA exclusively at the base of the sensillum (Supplementary Figure S1A), suggesting that nompA does not interact at the odor-receptor interface but rather acts as an extracellular scaffold or matrix component likely holding OSN dendrites in place. Subsequently, we co-stained OSNs and thecogen cells through use of the OR-coreceptor (Orco)-GAL4 and nompA-GAL4 drivers, respectively. Here, we observed hallmark features of thecogen cells, as described in morphological EM studies of *Drosophila* sensilla (Shanbhag et al., 2000), namely a closely and thinly OSN-enveloping cell, sheathing the inner dendrites, with a nucleus at the reported distance from the OSN, of an apposed, sheathing cell in close proximity with the OSN (Figure 1G). To confirm our expectation that nompA-GAL4 faithfully labels thecogen cells across a broad range of different types and morphological classes of sensilla, we asked whether this thecogen cell marking approach would label predicted locales. Specifically, we asked whether the marker would occur in ab3 sensilla, in different morphological types of sensilla, as well as whether they would occur in Orco-negative sensory neurons expressing IRs, such as those found in coeloconic sensilla of the sacculus. Indeed, thecogen cells marked with nompA co-occurred in all three cases: in ab3 (Or22a-immunopositive) sensilla (Figure 1H), in large basiconics negative for Or22a (i.e. ab1 and ab2) as well as trichoid sensilla characterized by their thick, rounded bases, termed basal drums (Shanbhag et al., 1999) (Figure 1I), and finally in Orco-negative coeloconic sensilla contained within the sacculus (Figure 1I and 1J). Interestingly, we also found thecogen cells in distal portions of the funiculus, without attributable Orco-positive neurons (Figure 1K, Supplementary Figure S1B). We also located the inverse case, of neurons without attributable thecogen cells (Figure 1L), as well as the proximomedial region of the funiculus opposite the arista where nompA-GAL4 seemed not to drive expression (Figure 1M), indicating that the nompA promoter is not ubiquitous nor universally active across the total set of thecogen cells of the funiculus. To address the previously suggested possibility that the promoter sequence of nompA-GAL4 is only transiently active during a narrow developmental window prior to eclosion (Larter et al., 2016), we checked whether reporter expression remained consistent between newly eclosed and older flies. The expression of the GFP reporter did not seem to be affected by age, as reporter expression was present in approximately equal amounts between antennae harvested from freshly eclosed flies and 5+ day old flies (Supplementary Figure S1C). In tandem, we also devised a fly line of genotype nompA-GAL4;UAS-GAL4;UAS-RFP which maintains constitutively active reporter expression, and observed no difference in staining between such flies and the more straightforward nompA>RFP genotype which may have been vulnerable to temporal downregulations in GAL4 expression (Supplementary Figure S1C).

We also surveyed the use of ASE5-GAL4 to mark tormogen cells by a similar immunofluorescence approach. We found a more widespread and uniform reporter expression across the entire funiculus (Figure 1N). ASE5-GAL4 marked the cytoplasm of more globular cells unalike to those of thecogen cells, indicating a marking of a different cell type (Figure 1O). To our knowledge, there have been no attempts to determine whether the GAL4 marker strictly marks tormogen cells, or perhaps whether it may in fact partially mark its sister trichogen cell. Using confocal imaging one cannot differentiate between the two, but we observe very even and confluent cellular tiling between the cells without overlap (Figure 1O). Given that tormogen and trichogen cells often fold and extend over each other (Shanbhag et al., 2000), this expected lack of overlap indicates that ASE5-GAL4 likely truly marks the tormogen cells of the funiculus. This is supported by the fact that such ASE5 promoters conjugated with lacZ and GFP reporters have been shown to mark tormogen but not trichogen cell types (Barolo et al., 2000).

Lastly, we attempted to compare and contrast available glial cell markers for the third antennal segment by use of the GAL4 expression system. We first drove the expression of nuclear GFP using the promoter of repo, a classical *Drosophila* glial homeodomain transcription factor expressed exclusively in glial cells, and observed the absence of any glial nuclei near sensilla (Figure 1P), confirming the attributable distinction between glial cells and the glia-like thecogen cells. Similarly, using a GH146-GAL4 line reported to stain antennal glial cells that migrate into the antenna during development (Sen et al., 2005), we observe cell bodies branching in manners reminiscent of axon-ensheathing Schwann cells (Figure 1Q). No signal could be observed near the sensilla/surface of the funiculus, suggesting that glial cells do not directly participate at the odor-interfacing periphery of the olfactory system, in line with expectations.

### A subset of tormogen cells respond during odor stimulation

It is not known whether tormogen cells respond during odor presentation, whether to odorants or to resultant OSN activity. To assess whether such responses exist, we used an open antennal preparation whereby a bisected portion of the antenna is exposed to liquid phase odor stimulation and imaged using fluorescence microscopy, as done previously (Mukunda et al., 2014, 2016; Miazzi et al., 2016; Halty-deLeon et al., 2018). We targeted the tormogen cell specifically using ASE5-GAL4 and expressed the widely used genetically-encoded indicator GCaMP6f as an intracellular probe for Ca^2+^ (Figure 2A). Surprisingly, we found an abrupt increase in intracellular Ca^2+^ following exposure of the antennal preparation to the odorant proxy VUAA1, a synthetic, non-competitive allosteric Orco agonist (Jones et al., 2011), with a complete absence of response to its solvent, DMSO (Figure 2B). Upon repetition, we additionally noticed in recordings that some tormogen cells were responding to VUAA1 pulses, while others were entirely non-responding. We reasoned that this may simply come as a result of not all sensilla being VUAA1-sensitive (e.g. Orco-negative sensilla), but perhaps also due to differential expression across age. Prior labeling attempts with ASE5-GAL4 have previously been suggested to be developmentally regulated and decrease in transcriptional activity with age (Larter et al., 2016). To test this possibility, we replicated the experiment with a batch of freshly eclosed flies (<12h after eclosion) and found decidedly fewer responders than with our typical batches of older flies of age 2-12 days (Supplementary Figure S2A), excluding the explanation that older flies experience deteriorating ASE5-GAL4 driven reporter expression with age.

**Figure 2.**
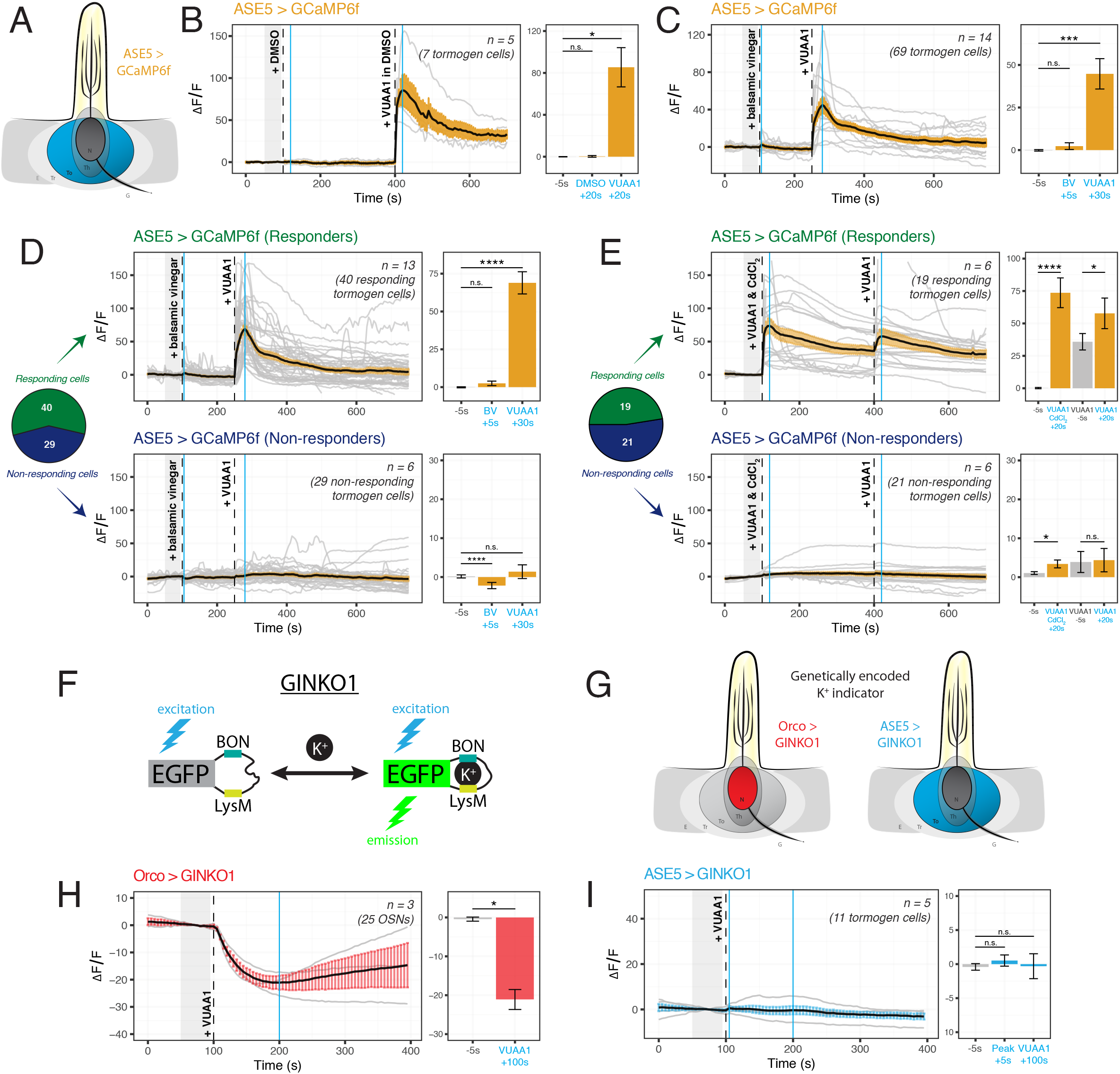
Cation dynamics in tormogen cells during odorant presentation. Tormogen cells are targeted using ASE5-GAL4. (**A**) Paradigm for Ca^2+^ imaging in tormogen cells. Calcium reporters are expressed specifically in the tormogen cell type using the GAL4/UAS system in an ASE5-driven manner. (**B**) Tormogen cells respond with an acute influx of Ca^2+^ following VUAA1 presentation. The effect is not attributable to solvent (DMSO) nor mechanical perturbations of liquid application to the imaged antennal sample. (**C**) Response of tormogen cells to presentations of 1:1000 balsamic vinegar and 100 μM VUAA1. Tormogen cells show no response to balsamic vinegar at this concentration. (**D**) Plot of Ca^2+^ imaging time course of individual cell responses (same experiment as shown in panel C), subsetted by response or lack thereof. A subpopulation (42%) of ASE5-GAL4 driven cells does not respond to odorant presentation. (**E**) Plot of CdCl_2_ voltage-gated Ca^2+^-channel-blocking experiment. VUAA1 in both presence and absence of 100 μM CdCl_2_ elicits Ca^2+^ influx into tormogen cells, but only in the responding subpopulation. Time courses and averages are plotted based on individual tormogen cell responses across all replicates. (**F**) VUAA1-responding and non-responding tormogen cells performed as seen in freshly eclosed flies. Time courses and averages are plotted based on individual cells. (**G**) Paradigm for K^+^ imaging in tormogen cells and OSNs. K^+^ cation reporter (GINKO1) is expressed using the GAL4/UAS system. (**H**) Olfactory sensory neurons exhibit an efflux of K^+^ following VUAA1 stimulation. (**I**) Tormogen cells exhibit no net K^+^ flux following VUAA1 stimulation. All error bars represent standard errors of the mean (SEM). Vertical blue lines indicate time points compared on each time course’s respective bar graphs. Paired student’s *t*-tests were used to compare time points. Asterisks indicate statistical significance (^n.s.^p>0.05, *p≤0.05, **p≤0.01, ***p≤0.001, ****p≤0.0001).

Given that tormogen cells are also present in sensilla equipped with OSNs that do not express Orco, such as those broadly found in coeloconic sensilla which instead express ionotropic receptors (IRs) sensitive to carboxylic acids and amines, we repeated the experiment with balsamic vinegar, which has been shown to elicit responses in IR-positive, Orco-negative OSNs (Jain et al., 2021). Moreover, we were prompted to test balsamic vinegar due to the fact that tormogen cells also occur in duplets in *Drosophila* coeloconic sensilla (Shanbhag et al., 2000; Nava Gonzales et al., 2021). Here we found no significant intracellular Ca^2+^ responses to vinegar as compared to VUAA1 stimulations (Figure 2C). Next, we qualitatively subset these cells into responding and non-responding groups for parallel analysis, based on whether Ca^2+^ rises were elicited upon either VUAA1 or vinegar stimulation during the experiment on a cell-by-cell basis. Thereafter we observed significant VUAA1-responding cells in the responding subgroup, and inversely, significant balsamic vinegar-responding cells within the non-responding subgroup (Figure 2D). Subsequently, using a similar approach to better understand the contribution of voltage-gated Ca^2+^ channels to the intracellular Ca^2+^ rise in tormogen cells, we used cadmium blocking of Ca^2+^ channels at a concentration of 100 μM CdCl_2_ (Wicher and Penzlin, 1997) and discovered that Ca^2+^ fluxes were evidently maintained in the presence of Cd^2+^-blocked Ca^2+^ channels within the responding subpopulation of tormogen cells (Figure 2E).

Next, we generated flies expressing the novel genetically-encoded K^+^ indicator GINKO1 (Shen et al., 2019) (Figure 2F) to determine whether OSNs, as well as tormogen cells, respond to odor presentation with intracellular K^+^ flux (Figure 2G). To validate the use of the indicator for the first time in *Drosophila*, we first tested the GINKO1 indicator driven by Orco-GAL4 expression in OSNs, as a test case where we would expect to see K^+^ efflux in OSNs during VUAA1 stimulation. Here we found a steady and substantial K^+^ efflux in OSNs (Figure 2H, Supplementary Figure S2B). Inversely, we found a complete lack of change or response to baseline K^+^ levels within tormogen cells following identical stimulations with VUAA1 (Figure 2I, Supplementary Figure S2C). Taken together, tormogen cells show strong Ca^2+^ influx upon odorant presentation, which does not seem to be dependent on Ca^2+^ channel ion flow, as well as no flux with respect to K^+^.

### Thecogen cells uptake K^+^ following VUAA1 stimulation

It is unknown whether thecogen cells, like tormogen cells, respond acutely during odor presentation. Using the same *ex vivo* antennal preparation and set up as in the previous experiment, we expressed the Ca^2+^ and K^+^ indicators CaMP6f and GINKO1 serially in the thecogen cells of the antenna using nompA-GAL4 (Figure 3A). Strikingly, and unlike the results in tormogen cells, we observe a complete lack of Ca^2+^ response to VUAA1 stimulation (Figure 3B). This is not the case however with thecogen intracellular K^+^ dynamics, which show a marginal increase following VUAA1 stimulation (Figure 3C). Because thecogen cells are relatively smaller in volume and size compared to other cells within the antenna (Shanbhag et al., 2000), and due to the small rise in intracellular K^+^ in thecogen cells following VUAA1 stimulation, we decided to validate the observed K^+^ rise and rule out the possibility of it being an experimental artefact. By lowering the concentration of K^+^ in the physiological medium five-fold, from 5 mM to 1 mM, we hypothesized that a K^+^ influx into thecogen cells should be enhanced following VUAA1 stimulation, as a result of increased K^+^ efflux into the extracellular lymph from neurons repolarizing at a lowered ambient K^+^ concentration (Contreras et al., 2021). Indeed, we observed a marked increase in peak response to VUAA1 stimulation in thecogen cells (Figure 3D). The results are suggestive of an excess extracellular K^+^ clearance mechanism, which has often been attributed to glial cells of the tripartite synapse as a homeostatic means to regulate the excitability of neurons, such as to prevent neuronal hyperexcitability (Walz, 2000; Sibille et al., 2015).

**Figure 3.**
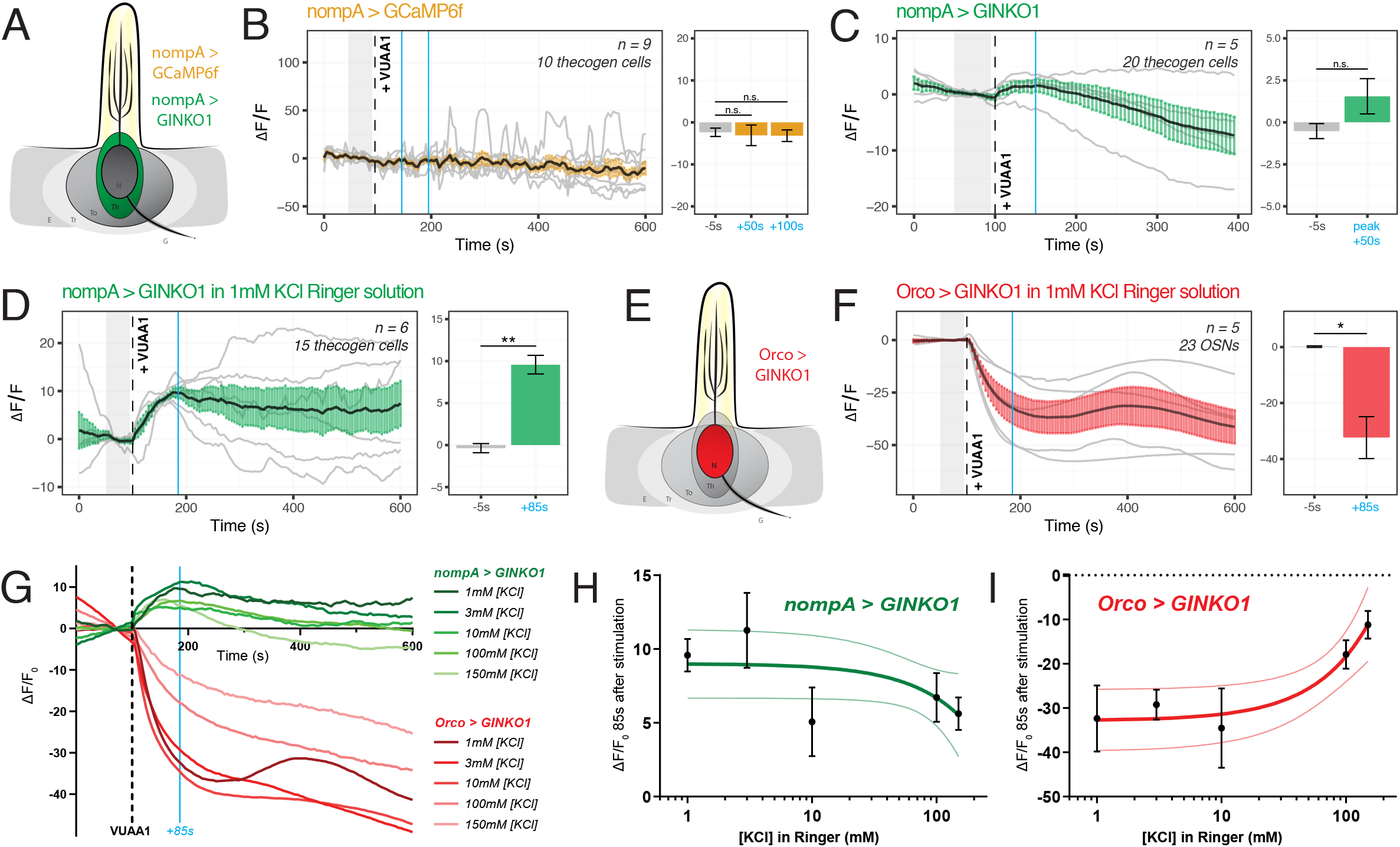
Cation dynamics in thecogen cells during odorant presentation. Thecogen cells are targeted using nompA-GAL4. (**A**) Imaging paradigm for Ca^2+^ and K^+^ in thecogen cells. Both Ca^2+^ and K^+^ cation reporters are expressed using the GAL4/UAS system in a nompA promoter-driven manner. (**B**) Live time course for Ca^2+^ imaging during VUAA1 stimulation. No net change in cytoplasmic Ca^2+^ in thecogen cells is evident following VUAA1 treatment. (**C**) Live time course for K^+^ imaging using GINKO1 during VUAA1 stimulation. A small increase in intracellular K^+^ concentration is observed. (**D**) K^+^ influx is enhanced following VUAA1 treatment at 1 mM KCl, a five-fold low extracellular concentration of K^+^ compared to panel C. (**E**) Illustration of K^+^ imaging in olfactory sensory neurons, as achieved by expressing GINKO1 in cells driven by the Orco promoter-GAL4. (**F**) K^+^ efflux is observed upon VUAA1 stimulation in olfactory sensory neurons. (**G**) Plot of average time courses across various extracellular KCl concentrations. Green: K^+^ imaging in thecogen cells, driven by nompA-GAL4. Red: K^+^ imaging in Orco-positive OSNs driven by Orco-GAL4. Cyan line indicates time point at which concentration curves are drawn in the following panel. (**H**) Concentration curve for responses at 85 seconds-post VUAA1 treatment in thecogen cells. (**I**) Concentration curve for responses at 85 seconds-post VUAA1 treatment in OSNs, indicating an inverse relationship in K^+^ ion concentrations between the two cells. All error bars represent standard errors of the mean (SEM). Vertical blue lines indicate time points compared on each time course’s respective bar graphs. Paired student’s *t*-tests were used to compare time points. Asterisks indicate statistical significance (^n.s.^p>0.05, *p≤0.05, **p≤0.01, ***p≤0.001, ****p≤0.0001).

To determine whether this explanation could hold, we looked at the effect of varying extracellular K^+^ concentrations on the intracellular K^+^ concentrations measured using the GINKO1 indicator expressed via the GAL4/UAS system (Figure 3E). First, we surveyed the neuronal dynamic with respect to K^+^ efflux at an extracellular [K^+^] of 1 mM, with the expectation of a larger efflux than with the conventional medium of 5 mM that approximates a physiological K^+^ concentration of the sensillum (Reinert et al., 2011) (see Figure 2H and Supplementary Figure S2B). As expected, we saw a longer and more prominent efflux of K^+^ at the relatively low extracellular [K^+^] of 1 mM (Figure 3F). We performed dose-response experiments for both OSN and thecogen cell with varying, physiologically relevant concentrations of K^+^, across 2 orders of magnitude between antennal- and sensillum-relevant concentrations of 1-150 mM (Reinert et al., 2011), and found a general, opposite flux pattern between OSNs and thecogen cells in peak K^+^ response, decreasing at higher ambient K^+^ concentrations (Figure 3G). A generally mirrored trend is observed upon plotting a dose-response curve at the average peak K^+^ influx time point of 85 seconds post-VUAA1 stimulation for thecogen cells (Figure 3H) and Orco-positive OSNs (Figure 3I).

In conclusion, we find that thecogen cells demonstrate no Ca^2+^ flux in response to antennae stimulated by VUAA1, in spite of concomitant OSN activation by VUAA1, but seem to respond to local extracellular rises in K^+^ by absorbing ambient K^+^ temporarily. These features are unlike the tormogen cell, which shows the exact opposite trend. The results with respect to cation movement within OSNs as well as thecogen cells are summarized in Table 2.

**Table 2.**
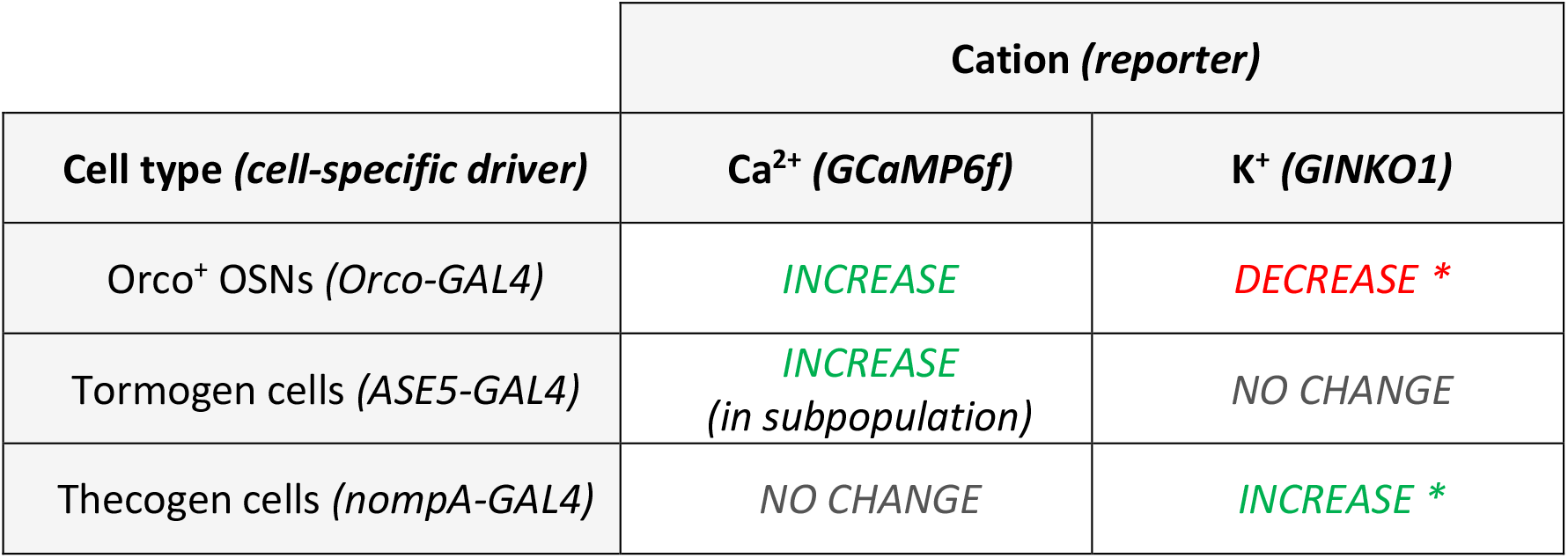
Summary of intracellular cation dynamics following odor presentation in neurons and support cells. Tormogen cells respond to VUAA1 application by influx of cytoplasmic Ca^2+^ while thecogen cells exhibit a cytoplasmic K^+^ influx. Asterisk (*) denotes a possible K^+^ buffering mechanism coupling OSNs and their thecogen cell.

### Thecogen cell ablation by induced apoptosis in adult flies affects neuron properties

Next, we asked whether the removal or absence of thecogen cells would have an effect on the response properties of neurons in sensilla *in vivo*. To achieve this, we generated flies with and without ablated thecogen cells. Because we were interested in adult olfactory properties, we chose to ablate thecogen cells after completion of eclosion and sensillum development (1+ day post-eclosion) to avoid developmental defects that occur in the case of an absence of support cells during sensillum biogenesis, thus avoiding confounding explanations such as improper development of the neuronal cilia or sensillum milieu (Chung et al., 2001; Andrés et al., 2014; Ando et al., 2019). Thecogen cell ablation by apoptosis was accomplished by targeting thecogen cells using the nompA-GAL4 construct, in tandem with GAL80^ts^ and UAS constructs. A repressing GAL80^ts^ construct incorporating a promoter of the ubiquitous housekeeping gene alpha-tubulin 84B (tub-GAL80^ts^) was used to reversibly inhibit the expression of the UAS-reaper (rpr) construct in targeted thecogen cells in a temperature-dependent manner. rpr, a caspase-dependent apoptosis-inducing protein (White et al., 1996) would thus be expressed upon heatshocking of flies at the GAL80^ts^-restrictive temperature of 32°C (Figure 4A). All flies were therefore reared at the GAL80^ts^-permissive temperature of 18°C from egg to eclosion to prevent rpr expression in thecogen cells. After eclosion, flies were separated into a control cohort kept at 18°C, and an ablation cohort that underwent 24h of heat-induced thecogen cell apoptosis at the GAL80^ts^-restrictive temperature of 32°C, wherein GAL80^ts^ loses its GAL4-repressive function and inhibition of rpr expression is lifted. The rearing and heatshocking schema for both experimental flies and parental controls is summarized in Figure 4B.

**Figure 4.**
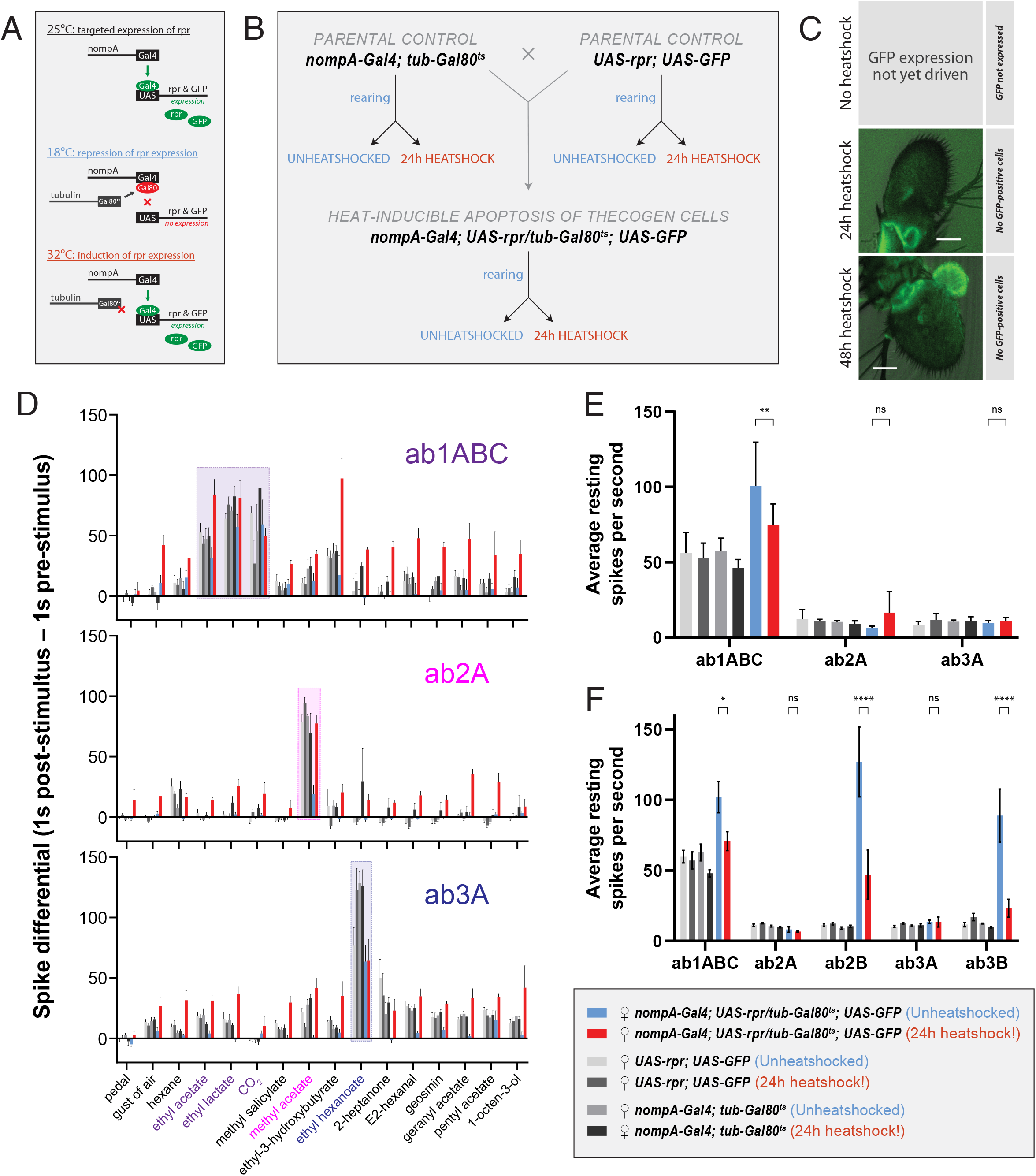
Effects of ablation of thecogen cells in adult flies by apoptosis on *in vivo* neuron responses to a panel of treatments. (**A**) Schematic of inducible ablation of thecogen cells by heatshocking using the GAL4/UAS system. (**B**) Crossing scheme used to generate control flies and thecogen cell-free adult flies. Experimental fly cohorts are split into heatshock and no heatshock conditions, as well as three genotypes, of which two are parental controls. (**C**) Confocal Z-stack of antennae following a 24h and 48h heatshock at 32°C, showing an absence of GFP signal, indicating complete absence and loss of thecogen cells following heat treatment. (**D**) Treatment response profile for 3 sensilla: a sum of spikes in all ab1 neurons excluding the D neuron, spikes of ab2A neuron, and spikes of ab3A neuron. Response profile is calculated by taking a sum total of all spikes 1s following stimulus onset subtracted from sum total of all spikes 1s prior to stimulus onset. (**E**) Average neuron activity per second measured from a 13s measurement window during no treatment. Units in resting spikes per second. (**F**) Average neuron activity per second measured from sum total of spikes preceding stimulus onset by 1s. Units in resting spikes per second. Two-way ANOVA with the Tukey *post hoc* tests were used to compare all sets of data; only some are shown. Asterisks indicate statistical significance (^ns^p>0.05, *p≤0.05, **p≤0.01, ***p≤0.001, ****p≤0.0001).

First, we checked whether heat induction would remove thecogen cells fully without trace. Here, we concurrently used a UAS-GFP construct to act as a label to follow the presence or absence of thecogen cells following induced apoptosis. We screened antenna to look for remaining GFP signal in the event of thecogen cells being leftover. Antennae were inspected closely, and no GFP signal across the entire depth of antenna was found following both 24h and 48h heatshocking periods during confocal imaging using long exposure and high excitation laser intensity, indicating a complete loss of thecogen cells (Figure 4C).

Next, to assess the neurophysiological properties of fly sensilla and neurons with and without intact thecogen cells, we used single sensillum recording (SSR) in female flies to screen the responses of 3 sensilla subtypes (ab1, ab2 and ab3) to a panel of 16 treatments (*n* = 4-9 female flies). As controls, we also tested both parental lines used to generate the experimental fly line where thecogen cell-specific apoptosis could be induced. We excluded the smallest neurons ab1D, ab2B and ab3B from most analysis due to poor signal-to-noise ratio rendering them difficult to uniquely identify among neighboring neurons and recording noise. First, we obtained a response profile to the panel of 16 treatments, including no treatment, a blank gust of air, and 14 ecologically-relevant odorants, of which some are diagnostic odors, i.e. best known ligands for the particular sensilla that were recorded from. For the gust of air and odorant presentation treatments, we used a 0.5 second stimulus duration. The concentrations of the odorants used are listed in the Materials & Methods section. A response profile for all treatments was calculated such that a count of all neuron spikes within 1 second prior to stimulus onset was subtracted from the count of all neuron spikes within 1 second following stimulus onset. In all fly groups, we observed generally conserved best ligand responses and unaltered odor tuning profiles with respect to tested sensillum or neuron odor responses, comparable to those in control flies (Figure 4D, Supplementary Figure S3A-B). However, we also observed broad non-specific responses in the heat-induced condition of experimental flies where thecogen cells were ablated. We reasoned that this could be explained by changes to the resting activity of the neurons, or that OSNs that had lost their thecogen cell support were gaining non-specific odor response profiles.

To address both possibilities, we noticed that blank gusts of air and hexane stimulations were eliciting responses in a manner restricted to only the experimental cohort of heat-shocked flies, most evident in ab1ABC and ab3A neurons (Supplementary Figure S3C). On this front, we mined the SSR data to estimate the resting activity of neurons prior to any treatment. Here, we took an average spike count per second on active SSR recording during the ‘no treatment’ recording (Figure 4E). We noticed a weakly significant decrease in resting activity in ab1ABC neurons between non-heatshocked and heatshocked flies, though evidently not different enough to those of baseline levels shared between parental and experimental cohorts. Moreover, there was no change in resting activity in ab2A and ab3A neurons, indicating that thecogen cell ablation has no effect on resting spontaneous activity in these neurons (Figure 4E). However, given that all three sensilla host multiple neurons, we additionally surveyed the resting activity of the small B neurons in the ab2 and ab3 sensilla which have been reported to be ephaptically coupled (Zhang et al., 2019). Here, we noticed a significant decrease with heatshocking, a phenomenon restricted only to the B neurons’ resting activities, though not in a meaningful manner relative to control flies (Figure 4F).

Finally, to understand the temporal nature of the neurophysiological responses as measured by SSR, we plotted response frequency traces for all odor presentations recorded in all flies and tested sensilla. We obtained average traces for recordings in ab1ABC, ab2A and ab3A neurons to 8 treatments: no treatment, a blank gust of air, the odor solvent hexane, and 5 odors: the ab1ABC diagnostic odorants ethyl acetate, ethyl lactate and CO_2_, the ab2A diagnostic odorant methyl acetate, and the ab3A diagnostic odorant ethyl hexanoate. Due to the observation of non-odor specific stimulation of neurons in thecogen cell-ablated flies, we reasoned that the neurons may have gained mechanosensitivity to odor presentations, given that all odor stimulations included air gusts. We attempted to dissociate the olfactory component of responses from the mechanosensitive response by analytically correcting all responses via subtracting an average blank air gust response on a per-sensillum basis (Supplementary Figure S4). Here, we wanted to verify whether any responses would remain that could be attributed to odor sensing, rather than the gain in mechanosensitivity resulting from thecogen cell ablation. Following a three-step analysis of all odor response traces in all cohorts, we found some remaining olfactory component in the ab1 sensillum, and no remaining olfactory component in ab2 and ab3 sensilla upon correcting traces for their mechanoresponse component (Supplementary Figure S5).

Aside from this, we also noticed several trends when comparing between heatshocked and unheatshocked flies, which were plotted using a bubble chart showing differences between heatshocked and unheatshocked cohorts. This was done for peak response frequency (Supplementary Figure S4C) as well as total response area-under-curve (Supplementary Figure S4D). We found no major effects restricted to odors specific to those sensilla between control flies with intact and ablated thecogen cells (dashed boxes, Supplementary Figure S4C-D). We additionally found no differences in neuron resting activity prior to stimulus onset, in line with our previous results (Figure 2E-F). However, we noted that the ab1 sensillum generally exhibits higher responses in thecogen cell ablated flies, while ab2 sensilla show no changes between heat treatment cohorts. Somewhat tentatively, for both peak frequency and area-under-curve comparisons, the ab3 sensillum exhibits lower responses in thecogen cell ablated flies in terms of area-under-curve as well as peak response frequencies (Supplementary Figure S4C-D), which may indicate different sensillum-specific tolerances of thecogen cell ablation. Nonetheless, from this data analysis we can only find tentative evidence for remaining olfactory sensitivity to odor presentations, whereby most of the contribution to treatment responses was found to come from gained mechanoresponses. We leave the data open to interpretation and as a reference for future studies.

## 4. Discussion

### Recap of study

In this study we have taken three broad approaches to begin to understand the role of *Drosophila* antennal support cells in odor perception. Initially, we set out to address the scarcity of systematic descriptions available prior to further investigation. First, we evaluated a variety of support cell type-specific genetic immunolabeling techniques to identify the validity, suitability and limitations of each as tools to target support cell types specifically, and characterized the cellular distribution of each across the funiculus. Of particular interest was the thecogen cell-specifying genetic driver nompA-GAL4, which we showed labeling thecogen cells across a variety of sensillum types, in both Orco-positive and Orco-negative sensilla, but which did not label the totality of all thecogen cells in the third antennal segment, a result unacknowledged in previous studies employing nompA promoters in chemosensory sensilla (Chung et al., 2001; Jeong et al., 2013). The observed absence of thecogen cell marking in the proximomedial region of the funiculus may only be explained in part by the zone’s enrichment with spinules, uninnervated and aporous hairs originating from epithelial cells, which are not underpinned by typical sensillum architecture (Shanbhag et al., 1999).

Second, by virtue of live cation imaging in an *ex vivo* preparation of antennal tissue, we found concomitant responses of tormogen and thecogen cells to OSN stimulations with the odor proxy VUAA1. We found that a subset of tormogen cells undergo an acute and steep cytoplasmic Ca^2+^ influx immediately following stimulation, without concomitant K^+^ influx, indicating the quick activation of tormogen cells during odor presentation events. The opposite trend was observed in thecogen cells, over a range of concentrations of ambient K^+^, pointing to the potential role of thecogen cells as ionic sinks involved in K^+^ buffering of the sensillum or perineuronal lymph.

Third, by way of removing thecogen cells in adult flies using inducible apoptosis, we assayed three distinct basiconic sensillum subtypes electrophysiologically using SSR for response profile changes following thecogen cell ablation. We firstly observed a broad loss of specificity to odorants, despite the generally conserved best-ligand property in thecogen cell-free sensilla in all three tested sensilla. We also noted the lack of any change in OSN resting activity following thecogen cell apoptosis, as well as in heatshocked controls. The broadly observed response to both mechanical air gust treatments and odorant pulses was attributable to a gain in mechanosensitivity, likely as a result of a loss of cellular or dendritic integrity within the sensillum architecture that depended on the thecogen sheath. This result is distinct from a previous report of odor insensitivity in nompA-null mutants, where sensillum biogenesis did not proceed correctly and where sensory dendrites were unable to innervate the sensillum (Chung et al., 2001). In our study, we induced the removal of thecogen cells at a post-development stage, and still found that tested neurons retained some degree of best ligand specificity, while observing a functional gain of mechanical responsiveness to gusts of air. These effects of thecogen cell ablation are in our view not attributable to the loss of support cell OBPs, given that our study has used comparable pulse durations and odor concentrations as previously in studies showing robust olfactory responses in the absence of basiconic OBPs (Xiao et al., 2019), and also that thecogen cells themselves have been shown only to express Obp28a in ab1-3 sensilla in a patchy manner (Larter et al., 2016). Hence, we posit that the effects of thecogen cell ablation are not attributable to the absence of any nascent OBPs which would otherwise exist in sensilla with intact support cells. Lastly, we attempted to remove the mechanoresponsive component of all responses, and found evidence of impaired odor sensing in OSNs, which becomes mostly but not entirely absent. We also tentatively suggest that thecogen cell loss may be tolerated differently by OSNs with respect to odor detection in a sensillum-specific fashion.

### Known and unknown mechanisms coupling sensory neurons and support cells

The coupling of support cells to OSNs remains enigmatic. Here we have demonstrated two separate kinds of response, of two distinct support cell types, to odor presentation events. This indicates a coupling in real time between the OSN and the supporting cells, and some yet unknown coupling mechanisms must exist to explain the observed interactions. How exactly do support cells sense the activation of their OSN, or perhaps even the arrival of an odor? Ostensibly, are OSNs interacting with neighboring cells by direct contact, perhaps through gap junctions, or communicating via paracrine signaling, or indirectly through their influence on the extracellular milieu, via ion exchange and homeostatic maintenance? The exact modes are currently unknown in *Drosophila* to date, though many lines of evidence may provide hints. Formerly, in some basiconic sensilla, we have found no evidence of gap junctions connecting neurons with support cells in experiments using gap junction-permeable dyes injected into the sensillum lumen that backfill the OSNs (unpublished data). However, one cannot exclude that support cells are directly connected between each other via gap junctions, or that a subset of sensilla may harbor neuron-support cell junctions and channels. Indeed, this may be a likely possibility given that the expression of insect gap junction proteins, namely the innexins ogre/inx1, inx2 and inx3, are expressed in substantial levels in adult fly antennae as surveyed by several independent studies employing antennal tissue RNA-seq (Menuz et al., 2014; Barish et al., 2017; Pan et al., 2017). Innexins in adult insect sensory systems have hitherto only been described in the auditory system, where they are involved in the synaptic coupling between auditory sensory neurons and the giant fiber (Pézier et al., 2016), but have been tentatively noted in early silk moth EM studies looking at membrane-to-membrane contacts (Steinbrecht, 1980). Gap junctions between sensory support cells have been more confidently described in the mammalian cochlea, where they actively function in cochlear amplification (Zhu et al., 2013). It remains an open question whether gap junctions may play a role in the olfactory periphery of *Drosophila* or other insects.

Another striking possibility is that supporting cells may detect odor cues directly. The ammonium transceptor Amt was recently reported to be widely expressed in *Drosophila* support cells along with a lesser population of ac1 neurons (Vulpe et al., 2021). If this is not incidental, we propose the possibility that the wide Amt expression in support cells of coeloconic sensilla could allow for some degree of enhanced ammonia odor sensing or processing. This could have an indirect function in potentiating ac1 neurons, such as with *C. elegans* glial cells detecting odorants separately from sensory neurons allowing for support cell-dependent adaptation of neuronal responses (Duan et al., 2020), especially in the ecological context of environments where relevant, common cues such as ammonia can be present in excess. A similar phenomenon may also exist for tastants probed by *Drosophila* gustatory sensors, which also harbor Amt transceptors (Delventhal et al., 2017).

On the other hand, modes of paracrine signaling between support cells and sensory neurons have been reported in other sensory systems. In sensilla contained within the amphid of *C. elegans*, amphid sheath glial cells have been shown to suppress sensory neuron activity and promote chemosensory adaptation through local release of GABA (Duan et al., 2020). More generally, *C. elegans* glia have long been known to tune sensory neuron activity, which in turn affect behavioral responses to environmental stimuli (Wang et al., 2008, 2012; Han et al., 2013; Stout et al., 2014). Yet another case of paracrine signaling is within the mouse cochlea, where supporting cells adjacent to mechanoreceptive neurons express the ATP-sensitive purinergic membrane receptor P2RY1. When activated, the support cells experience a cytoplasmic Ca^2+^ influx from intracellular stores, followed by K^+^ efflux from the support cells which in turn elicit local neuronal burst firing (Babola et al., 2020). Interestingly, the release of K^+^ in this model leads to a rapid increase in extracellular space due to support cell crenation, which may have consequences in subsequent K^+^ redistribution and the termination of neuronal firing (Babola et al., 2020). Though this is a case study from an immature and developing sensory system, it may be an indication of potential mechanisms for how support cells may influence sensory neurons by purely ionic and physical responses within a contained compartment such as that of the sensillum lymph, by means of swelling and shrinking and modulating the ionic milieu of the environment. For one, the morphological shape of *Drosophila* supporting cells, with respect to the aqueous lymph, seems to freely allow such mechanisms. Indeed, swelling has been suggested in older studies where moth thecogen cells have been observed to manifest signs of swelling with increasing incubation time with lanthanum ion solutions, as well as unusually close apposition of thecogen and neuronal membranes following freeze substitution (Steinbrecht, 1980; Keil and Steinbrecht, 1987), which may provisionally indicate some intrinsic capacity of support cells to deform lymph spaces.

More yet, in the mouse olfactory epithelium, it has also been demonstrated that olfactory ensheathing cells (OECs) are sensitive to neighboring neuronal activity, as observed in cytosolic increases of Ca^2+^, and that such coupling is dependent on ATP and glutamate release from the neuron (Rieger et al., 2007). Further evidence of purinergic junctions between neurons and non-neuronal cells have also been implicated in microglia within their role in regulating neuronal Ca^2+^ load, and both monitoring and protecting neuronal function through specialized ultrastructures (Cserép et al., 2020). Whether similar modes of paracrine cell-cell signaling occur in *Drosophila*, coupling the activity status of OSNs to support cells, remains an open question. Such considerations may help broaden our view of odor processing in insect peripheral sensory tissues and aid in generating new, non-canonical hypotheses.

### Ion homeostasis and indirect effects

Release and uptake of K^+^ is entangled with several dynamic processes, namely the generation of action potentials and dendritic potentials, as well as the further concentration and release of K^+^ from the same or neighboring cells (Ransom et al., 1986; Syková, 1991; Ransom, 1992; Ransom and Ye, 1995; Walz, 2000). Though difficult to compare in our data due to differing shapes and genetically-encoded cation indicators used, tormogen cells seem to exhibit a much quicker Ca^2+^ response than the K^+^ sequestering occurring in thecogen cells, perhaps due to aforementioned changes in shape or due to quicker mobilization of intracellular Ca^2+^ stores, which may hint at the active role of tormogen cells in maintaining properties of the sensillum lymph space through mechanisms such as export of accessory proteins. And though often overlooked, the possible existence of an isolated perineuronal cleft between the thecogen cell and OSNs in *Drosophila* must also be taken into regard, which may differ in ionic composition from the sensillum lymph, a notable feature described in moths (Steinbrecht, 1980; Keil and Steinbrecht, 1987). At least in some moth species, the thecogen cell and neuron have been purported to contribute jointly to the electrical properties of the sensillum (De Kramer et al., 1984; De Kramer, 1985; Kaissling, 1986).

Moreover, it is worthy of note that the glia-like thecogen cells are reminiscent of glial cells of the tripartite synapse model, where glia are thought to modulate synaptic communication through K^+^ ion buffering, dispersion and redistribution (Beckner, 2020). This is not a novel comparison. To some degree, chemosensory organs like sensilla have unmistakable similarities with neuronal synapses, and have thus been compared and used to model synaptic clefts, which also feature arrangements of receiving neurons and non-neuronal players which evolve together to respond to small, transient chemical signals (Shaham, 2010). In the same vein, perhaps we can also apply concepts from the role of the glial cell in the tripartite synapse to chemosensory systems in efforts to generate novel hypotheses. For instance, does the thecogen cell share features with the astroglial cradle (Nedergaard and Verkhratsky, 2012), shielding the neuron from the known multitude of external factors such as an ever-changing sensillum lymph chockful with accessory proteins and debris? Are ensheathing thecogen cells perhaps playing an essential role in sensilla that are innervated with multiple neurons experiencing interneuron ephaptic inhibition (Zhang et al., 2019)? Furthermore, recent evidence in *C. elegans* has shown that peripheral glial support cells engulf, phagocytose and prune thermosensory neuronal endings depending on their activity load (Raiders et al., 2021). Upon disruption of this pruning, a behavioral temperature preference is lost (Raiders et al., 2021). Do similar processes exist in insect sensillum repertoires? This is stated in light of the fact that compartmentalization of sensory neurons has independently evolved many times, which act as means to integrate sensory inputs at the earliest stages of odor processing (Ng et al., 2020). However, neither presence nor potential contributions of support cells are stressed, even though they may be foundational.

The K^+^ concentration ranges of insect antennae are also particularly notable. A particle-induced X-ray emission study of the *Drosophila* antenna indicates that the [K^+^] at the center of the funiculus is 50 mM, while at the sensillum edges it is 5 mM, a 10-fold lower concentration (Reinert et al., 2011). At face value, these results can be interpreted as vastly differing ion concentrations between the hemolymph and sensillum lymph. The difference may be explained by separately maintained, isolated compartments, physically sheltered from the OSN by the triad of support cells, in a manner similar to the perilymph and endolymph of mammalian cochlea (Zdebik et al., 2009). Analogously, sensillum lymph space as well as the perineuronal lumen between thecogen and OSN cells would be maintained separately from the hemolymph, and would thus exhibit drastically different ambient [K^+^] by virtue of enclosure or by concentrating K^+^ from the surroundings. Counterintuitively, sensillum lymph in insects has been reported to be high in K^+^ (Thurm and Kuppers, 1980; Steinbrecht, 1989) alike to the vertebrate endolymph (Corey and Hudspeth, 1979). We speculate that similar principles may apply to the olfactory sensilla of *Drosophila*, namely that the perineural cleft between thecogen and OSN are functionally distinct from the sensillum lymph, an idea long suggested (Keil and Steinbrecht, 1987). Tangential support for such a hypothesis has been observed in the nearby *Drosophila* auditory system, within the Johnston’s organ of the second antennal segment. Scolopale cells, a cell type homologous to that of thecogen cells, have been found to express the Na^+^/K^+^ ATPase pump preferentially localized on the side facing the perineuronal lumen, and have been suggested to pump K^+^ ions to maintain a K^+^-rich ionic presence for the sensory neuron (Roy et al., 2013). It is reasoned that sensory transduction events deplete K^+^ and thus require active replenishment. The knockdown of an ion pump subunit via RNA interference hence resulted in deafness, loss of scolopale cell integrity, and additional morphological defects such as the presence of swollen cilia implying an ionic imbalance in the scolopale space (Roy et al., 2013). In our experiments we similarly observe the loss of cellular integrity following thecogen cell ablation, with similar sensory detriments to odor detection, namely a loss of specificity to odorants, and speculate that similar explanations may at least partially account for the observed olfactory dysfunction.

### Future perspectives

A tremendous breadth of questions remains to be answered. Functionally, do *Drosophila* support cell activities influence classic neuronal properties such as adaptation and sensitization (Wicher and Miazzi, 2021), for example in their ability to discriminate transient, repeated, sustained or excessive cues, perhaps by switching rates of odor clearance from the sensillum lymph through adjustable release of enzymes or pinocytosis? Do support cells vary heterogeneously and phenotypically between sensilla within or between organisms, beyond that of known differential expression of OBPs? Curiously, to our knowledge a single study exists where support cell heterogeneity has been found: thecogen cells in moths are relatively enlarged particularly in hygro- and thermo-sensitive sensilla (Steinbrecht et al., 1989). Though we show responding and non-responding subpopulations of tormogen cells in this study, further diversifying the class of auxiliary cells, we hypothesize that many other support cell heterogeneities exist.

By the same token, do support cells found within larval olfactory systems (Hartenstein, 1988) share similar principles with those of the imago? What are the functions of the numerous extracellular vesicles observed in *Drosophila* sensilla, suggested to arise from olfactory support cells (Nava Gonzales et al., 2021)? Are various support cell functions conserved across sensory modalities, in other structures such as chordotonal organs, and in sensillum classes that serve gustation and hygro- or thermo-sensing? Do support cells have any roles to play in the neuronal turnover in insects (Fernández-Hernández et al., 2020) as they do in the adult mouse vestibular system (Bucks et al., 2017) or the avian inner ear (Bird et al., 2010)? With the advent of next-generation single-cell sequencing techniques, and with promising advances of applying these approaches to *Drosophila* antennae (unpublished data, McLaughlin et al., 2021), we may soon have access to the transcriptomic landscape of the antenna on a cell-by-cell basis to better understand the distinct varieties of both neuronal and auxiliary cells. Practically, we may get a glimpse into the set of underlying molecular players such as junction proteins, purinergic receptors and ion channels, and to which tissues, sensilla, modalities, sexes, internal states and life stages of the fly these can be attributed. Future research must take into account percipient evidence of the varieties of response modes between support cell types. Namely, we theorize that Ca^2+^ flux as observed in tormogen cells may relate to yet unknown (intra)cellular signaling processes, while K^+^ flux specific to thecogen cells may relate to homeostatic feedback mechanisms that may be subject to modulation (Walz, 2000).

Last, we may one day answer the question of whether support cells experience natural variation, and whether they can be a locus for selection in evolution in light of olfactory performance. It may be that support cells evolve changes to supplement the sensory periphery, just as neurons have (Prieto-Godino et al., 2017, 2020). We believe a synthesis of knowledge between the neuroscience of glial cells and sensory biology will cultivate a new appreciation for the complexity and functional interplay between both neuronal and supporting cells of sensory organs, and illuminate new mechanisms underlying physicochemical perception.

## Author contributions

SP, SLL and DW conceptualized the study. SP performed all immunofluorescence microscopy, fly rearing and crossing, cation imaging, formal analysis and visualizations. VPM performed single-sensillum recordings. SP and VPM processed the single-sensillum recordings data. SP and VV wrote the *R* script for automated processing and plotting of cation imaging data, as well as the script for the three-step analysis of the electrophysiological data and the script extracting the peaks and area-under-curve data from said SSR frequency traces. SP wrote the manuscript and prepared all figures. SP was supervised by DW and SLL. DW and SLL provided comments and critical input for manuscript revisions. DW and BH provided experimental resources. All authors have read and agreed to the published version of the manuscript.

## Acknowledgements

The authors thank Sabine Kaltofen, Regina Stieber-Rödiger, Silke Trautheim, Dr. Veit Grabe and Carolin Hoyer for technical lab assistance. We also thank Dr. Yi Shen for providing the GINKO1 plasmids, Dr. Fabio Miazzi for preparing the pUAST-GINKO1 construct for fly transformation, and Dr. Craig Montell for supplying the ASE5-GAL4 and nompA-GAL4 fly lines.

## Data Availability Statement

All raw and processed datasets and materials used in this study can be found in EDMOND, the data repository of the Max Planck Society, at the following location: https://dx.doi.org/10.17617/3.7m

## Figures

The figures appear below. Supplementary figures are available in a separate file.

