## Supplementary Figures for "Functional interaction between *Drosophila* olfactory sensory neurons and their support cells"

### Supplementary Figure S1

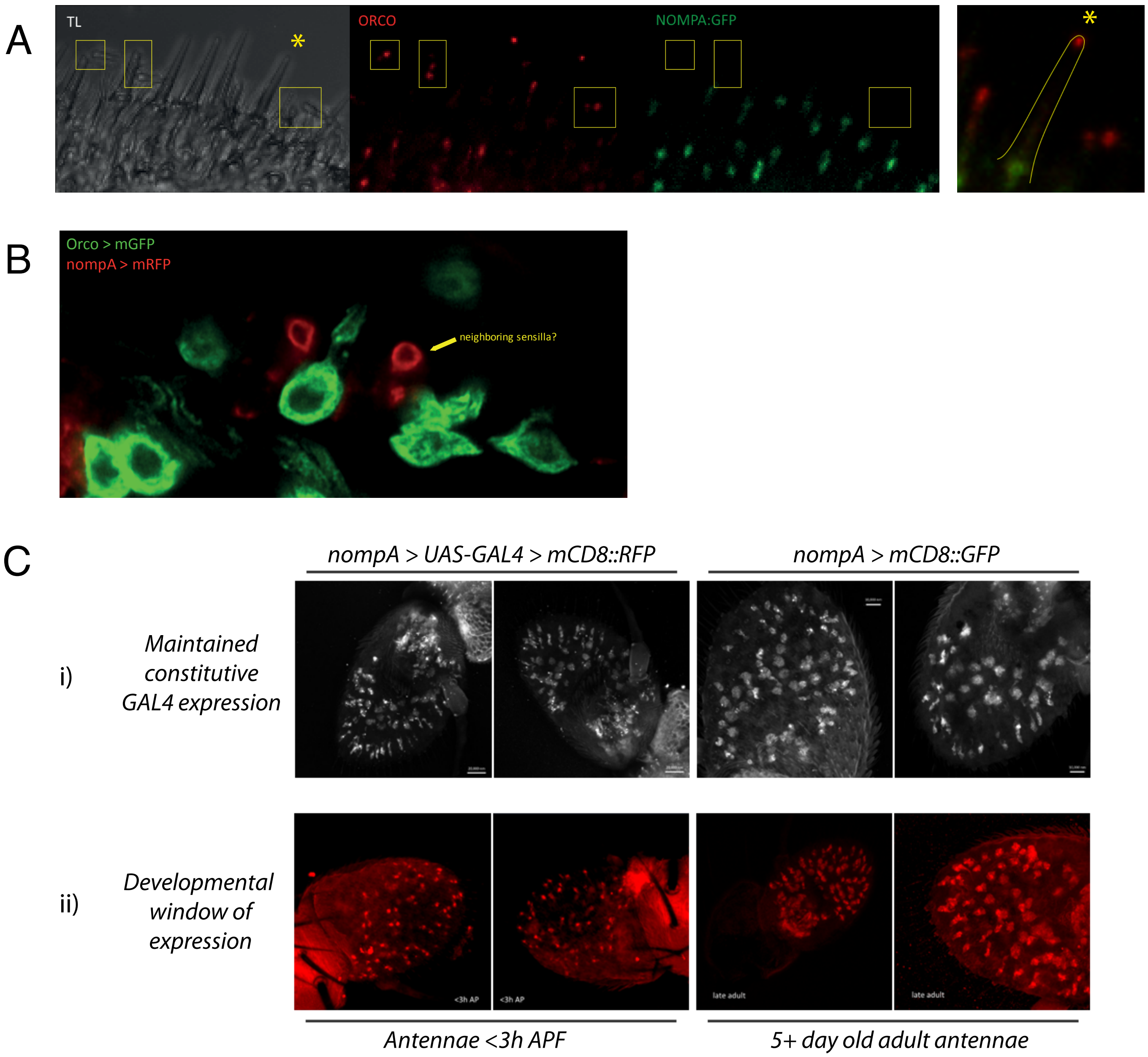

**Supplementary Figure S1. Addendum to Figure 1.** (**A**) Close-up of confocal imaging of Orco>RFP, nompA::GFP flies, indicating the sensillum-basal localization of nompA protein. (**B**) Close-up of portion of funiculus with two clear sheaths labeled with nompA>RFP, within which no Orco-positive OSN dendrites are found. (**C**) Comparison of labeling to check for developmental variation in nompA-GAL4 driven reporter expression. We compared regular staining (nompA>GFP, upper right panels, n=2) with maintained constitutive GAL4 expression (nompA>UAS-GAL4>RFP, upper left panels, n=2) and find no qualitative difference in expression between them. Likewise, we compared the fluorescent reporter staining of antennae of old flies (5+ day old flies, bottom right panels, n=2) with those of freshly eclosed flies (<3h after eclosion, bottom left panels, n=2) and similarly found no difference in expression levels, indicating that age does not play a decisive role in reporter expression across age.

### Supplementary figure S2

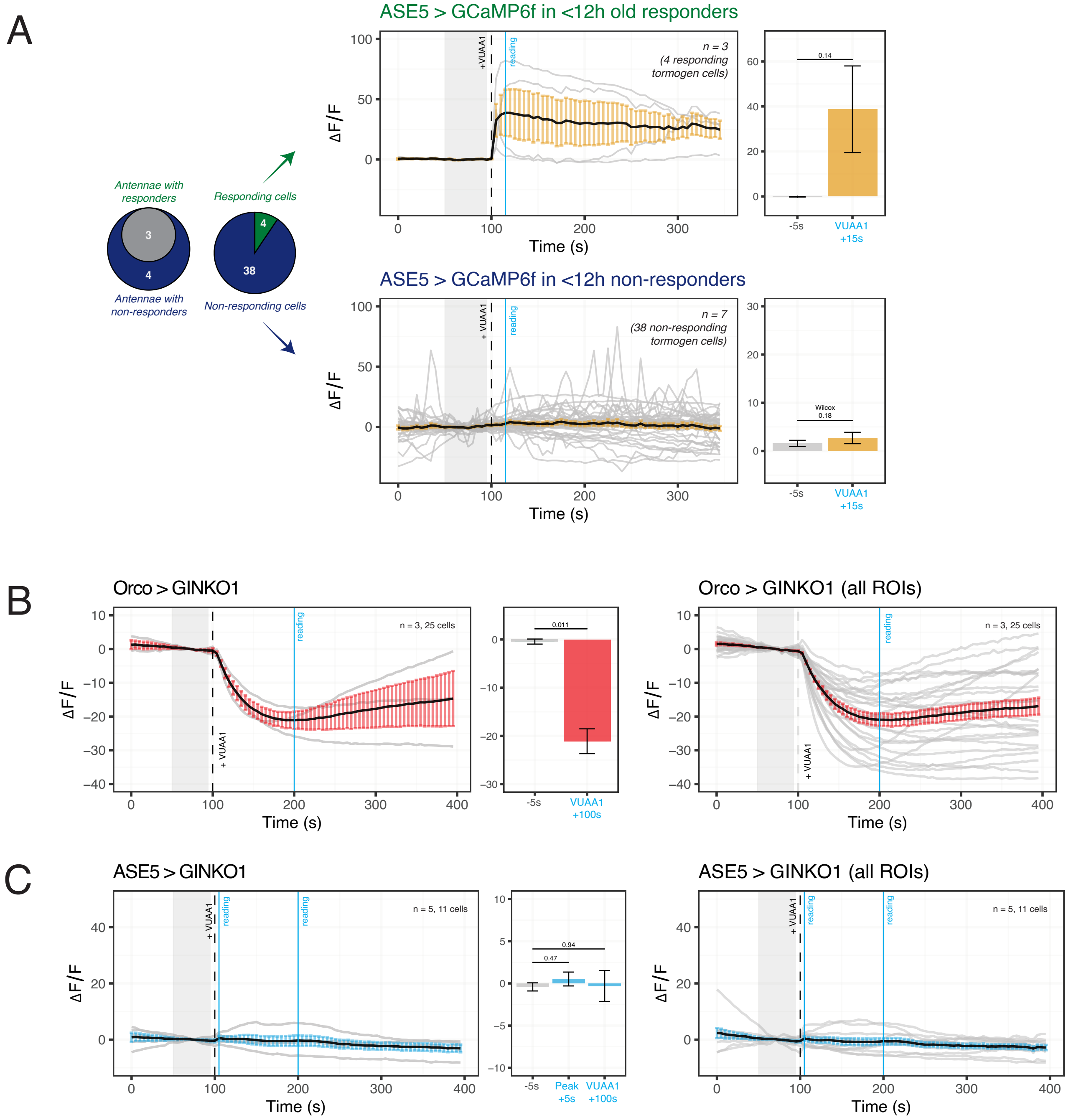

**Supplementary Figure S2. Addendum to Figure 2.** (**A**) $Ca^{2+}$ imaging of tormogen cells targeted using ASE5-GAL4, subjected to a stimulation of VUAA1 in an open antennal preparation from flies immediately after eclosion (<12 hours post-eclosion). There is a smaller proportion of responders in total than from older flies (see Figure 2D) but responders still maintain responsiveness ($Ca^{2+}$ influx) upon VUAA1 stimulation of the antenna. (**B**) Expansion of Figure 2H showing $Ca^{2+}$ imaging time course averages in OSNs targeted using Orco-GAL4, displayed per animal (left) as well as per all OSNs ("all ROIs"). (**C**) Expansion of Figure 2I showing $Ca^{2+}$ imaging time course averages in tormogen cells targeted using ASE5-GAL4, displayed per animal (left) as well as per all tormogen cells ("all ROIs").

### Supplementary figure S3

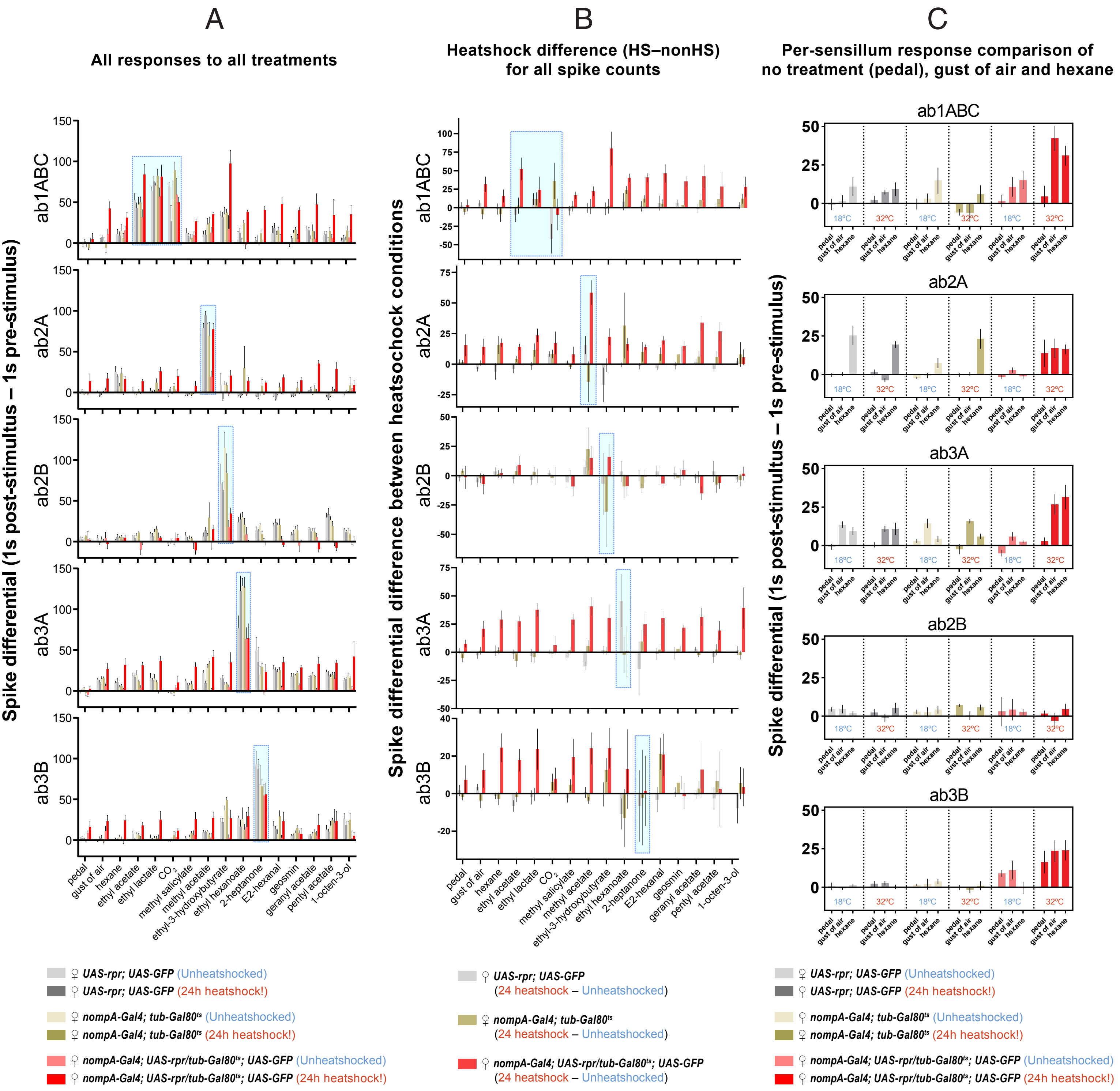

**Supplementary Figure S3. Addendum to Figure 4.** Small B neurons of ab2 and ab3 sensilla are additionally shown. (**A**) The responses to all treatments as shown in Figure 4D, but including the responses of the small B neurons in the ab2 and ab3 sensilla. Diagnostic odorant responses are highlighted within blue boxes. (**B**) Plot of the difference between heatshock and non-heatshocked fly cohorts of the same genotype (see legend). Diagnostic odorant responses are highlighted within boxes. Two-way ANOVA with the Tukey *post hoc* tests were used to compare all sets of data (not shown). (**C**) Per sensillum response to three SSR 'control' treatments, including no treatment ("pedal"), a blank gust of air, and hexane, the solvent used for the odorants. The data is the same as shown in Figure 4D.

### Supplementary figure S4

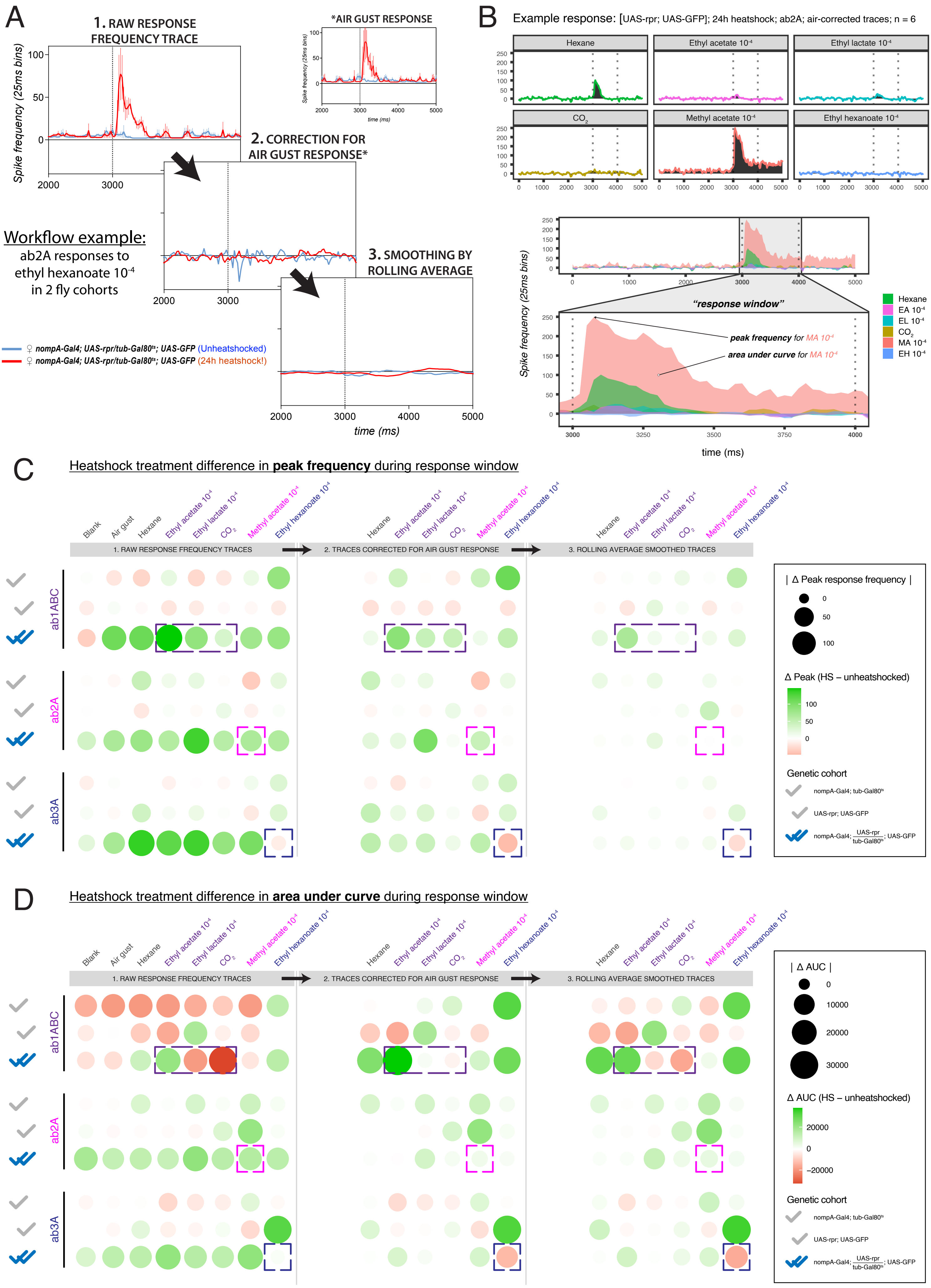

**Supplementary Figure S4. Three-step quantitative analysis of temporal time courses of responses of 3 sensilla to diagnostic odorants and treatments**. (**A**) Example of analysis pipeline for response of ab2A to ethyl hexanoate stimulation. All responses are plotted as a frequency time course. The first step shows raw response traces, as measured by SSR. A second step consists of correcting the response for the electrophysiological response to a gust of air by subtracting the average response to an air gust stimulus (*) at each time point for that sensillum. A third step consists of a 400 ms rolling average smoothing to eliminate leftover response artefacts caused by micro-timing mismatches between stimulus onset arrivals. We selected a 400 ms duration for smoothing because the bulk of most responses were of approximately 400 ms long durations (Supplementary Figure S5), and would thus constitute a conservative smoothing window. This analysis was performed for all 3 sensilla across 3 fly cohorts, 2 heat treatment conditions, for all 16 treatments. (**B**) An example time course for how peak response and area-under-curve during the 'response window' are determined. The response window is defined as 1s following stimulus onset, except for the third smoothed data analysis step, which is defined as 0.5 seconds prior to stimulus onset to 1 second after stimulus onset, due to shifting of response timing which occurs as a result of smoothing. (**C**) Bubble chart showing differences in peak frequency responses between heatshock treatment differences (24h heatshocked flies minus unheatshocked flies) for all 3 analysis steps. Dashed boxes indicate responses in thecogen cell-ablated cohorts for diagnostic odorants particular to that sensillum; following the three-step analysis, ab1 sensillum shows higher responses in thecogen cell-ablated (heatshocked) flies, ab2 sensillum shows no change, and ab3 sensillum shows lower responses in ablated flies. (**D**) Bubble chart showing differences in response area-under-curve during the response window, between heatshock treatment cohorts (24h heatshock-treated flies minus untreated flies) for all 3 analysis steps. Dashed boxes indicate responses in thecogen cell-ablated cohorts for diagnostic odorants particular to that sensillum; the results generally coincide with those in the previous panel, comparing changes in peak frequency of responses.

### Supplementary figure S5 (i)

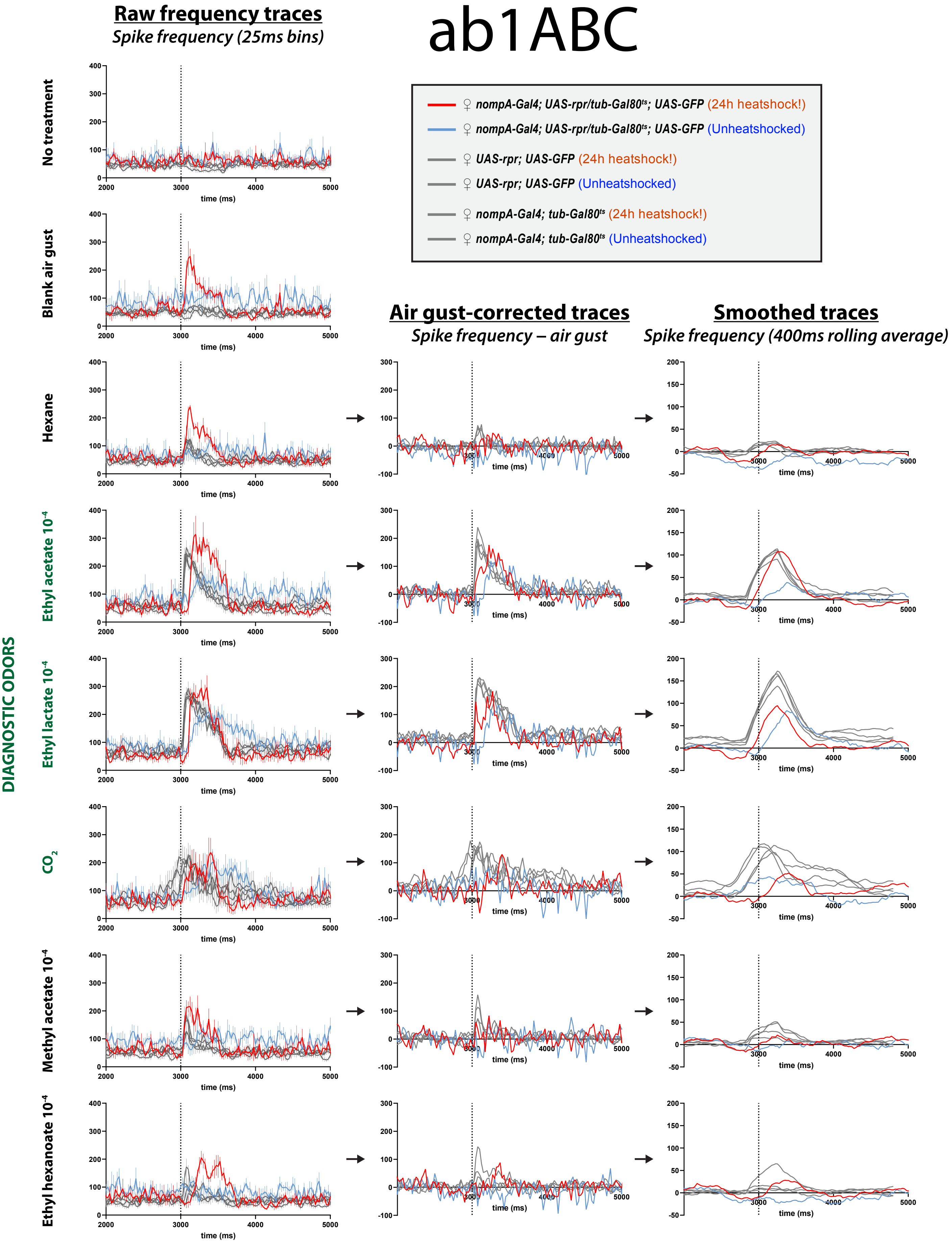

### Supplementary figure S5 (ii)

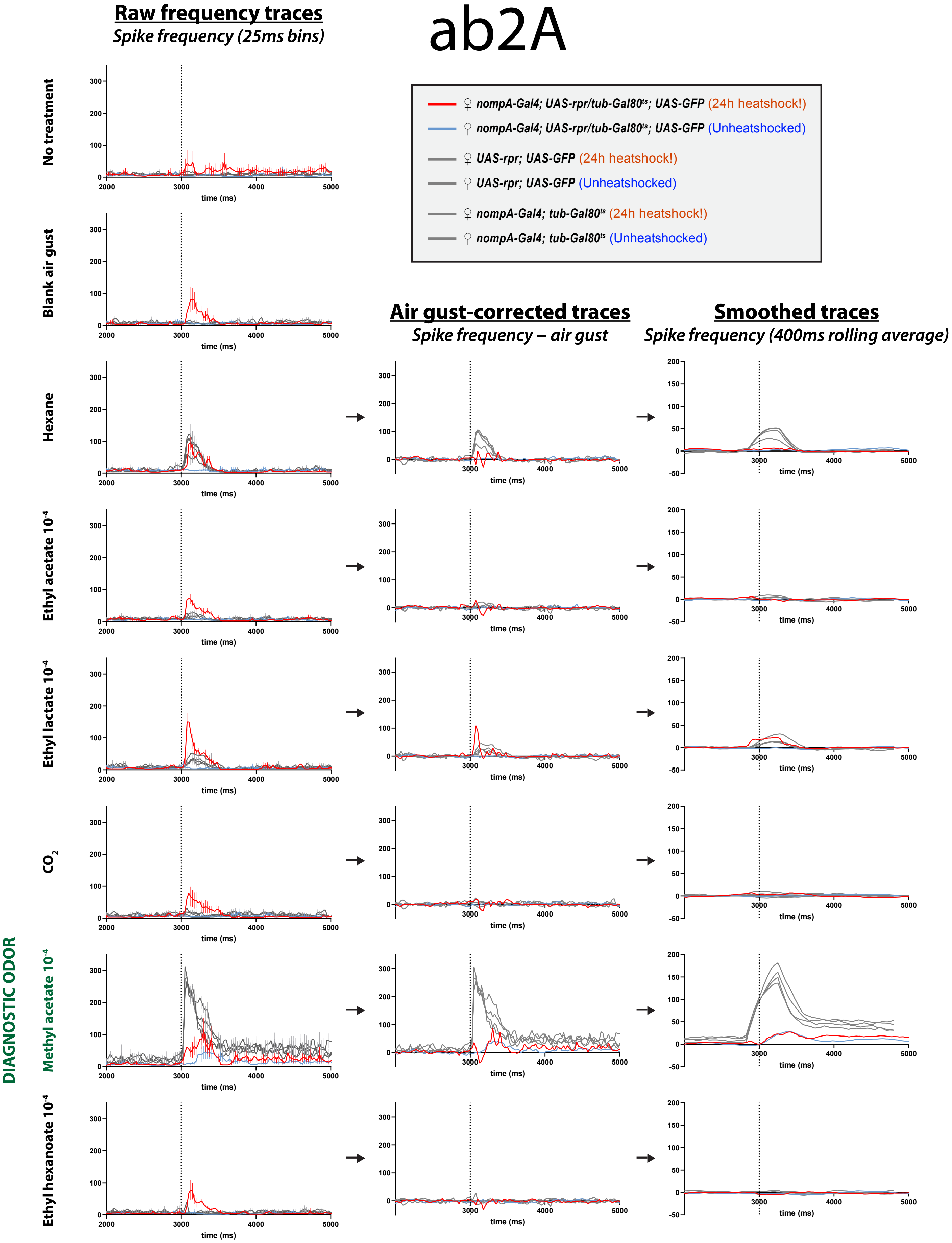

### Supplementary figure S5 (iii)

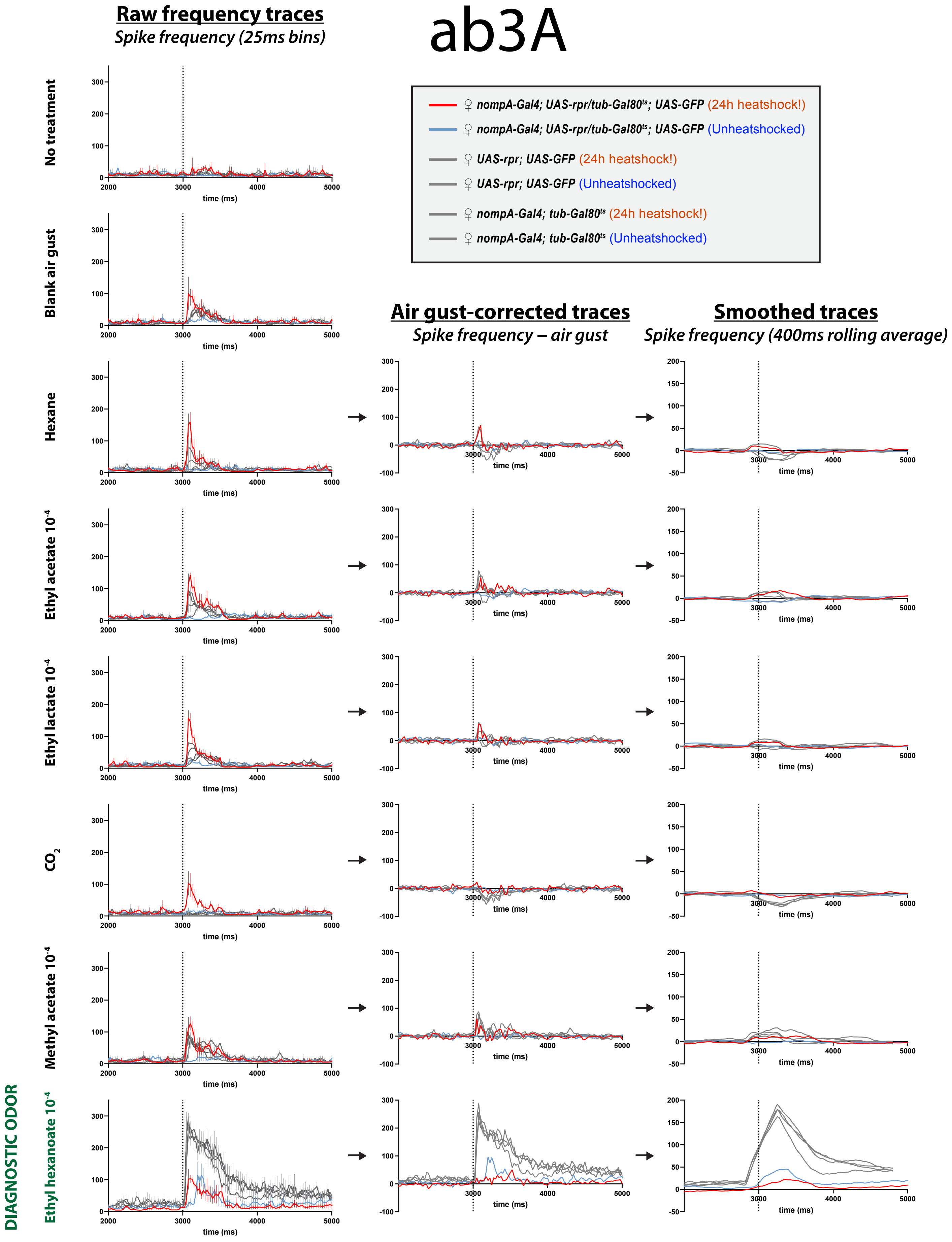

**Supplementary Figure S5. Raw and mechanoresponse-corrected response frequency traces in 3 sensilla.** A summary of the SSR recording data of responses in sensilla (**i**) ab1ABC, (**ii**) ab2A and (**iii**) ab3A, across all 3 data processing steps: first (left) column shows raw frequency traces of responses to odorants, second (middle) column shows air gust-corrected traces where responses to odor presentation gusts of air are removed from all responses to odorant pulses, and third (right) column shows the air-gust corrected traces smoothed by a 400 ms rolling average for that specific odor pulse. The smoothing third step was performed to remove artefacts caused by micro-timing mismatches between treatment pulse arrivals; 400 ms was selected because the bulk of most responses were of approximately 400 ms long durations, thus constituting a conservative curve-smoothing window. An example of the workflow can be seen in Supplementary Figure 4A. Diagnostic odor for the particular sensillum is highlighted in green.
